# Continuous and discrete brain dynamics to study behavioral adaptation during cognitive–motor dual-tasking in younger and older adults

**DOI:** 10.64898/2026.08.04.742700

**Authors:** Yan Deng, Daniel Kristanto

## Abstract

Dual-task paradigms are widely used to detect age-related cognitive and motor decline. Conventional evaluations typically average performance across an entire dual-task condition and compare it with a single-task baseline, implicitly treating performance as stable throughout testing. We challenged this assumption by examining block-wise behavioral adaptation and its dynamic functional-connectivity correlates. Forty older adults (50–80 years) and 20 younger adults (20–40 years) performed a cognitive Go/NoGo task, a motor pedaling task, and a combined cognitive–motor dual task during functional magnetic resonance imaging (fMRI) using a custom-built MRI-compatible pedaling device. Motor reaction-time (RT) variability was assessed across eight dual-task blocks, and dual-task benefit was defined as the relative reduction in variability from the first to the final block. Dynamic functional connectivity was characterized using two complementary approaches: dynamic independent component analysis (dyn-ICA), capturing continuously varying circuit properties, and a hidden Markov model (HMM), identifying recurring discrete network states.

Across participants, motor RT variability was highest in the first dual-task block, progressively decreased to its lowest level at Block 6, and remained comparatively stable thereafter. Both age groups achieved behavioral stabilization but followed distinct trajectories. Older adults progressed from pronounced initial variability toward their single-motor reference while continuing to perform the dual task, whereas younger adults began closer to this reference and maintained comparatively stable performance.

Greater dual-task benefit was associated with higher mean strength of a broadly distributed dyn-ICA circuit encompassing attentional, control, sensorimotor, visual, cerebellar, and default-mode systems (Circuit 4), as well as greater temporal variability of a functionally distinct circuit (Circuit 2).HMM analyses similarly linked greater benefit to more frequent visits to State 6 and greater occupancy of State 9, configurations involving coordinated sensorimotor, salience, dorsal-attention, and frontoparietal systems. Across both approaches, network features preferentially expressed by older adults were associated with greater relative benefit, whereas younger-enriched features accompanied smaller changes from a more stable initial level.These findings demonstrate that dual-task performance evolves substantially within a single session and that behavioral stabilization is related to both continuous circuit properties and discrete network-state visitation. Healthy older and younger adults may therefore achieve successful cognitive–motor adaptation through distinct regimes of dynamic whole-brain organization.

## 1. Introduction

Age-related deterioration in gait control is often expressed as increased gait variability, reflecting reduced stability and consistency of motor performance (Balasubramanian et al., 2015; Brach et al., 2007; Kowalski et al., 2022). When walking is combined with a cognitive task, this vulnerability may become more pronounced, producing elevated cognitive–motor dual-task costs (Lino et al., 2023; McPhee et al., 2022). Increased gait variability and elevated cognitive–motor dual-task costs are among the early behavioral markers of cognitive decline, incident dementia (Kwak et al., 2023; Montero-Odasso et al., 2017; Q. Yang et al., 2020), and Parkinson’s disease (Kim et al., 2024; Longhurst et al., 2023; Zhang et al., 2022). Cognitive–motor dual-task paradigms have therefore become an important framework for investigating how aging alters interaction between motor and executive control systems (Marusic et al., 2019; Piche et al., 2023). A prevailing account proposes that motor behavior becomes less automatic with advancing age and increasingly relies on executive control processes (Clark, 2015; Kvist et al., 2026; Montero-Odasso et al., 2012). Thus, although older adults may maintain relatively preserved performance during simple motor tasks, the addition of cognitive demands can expose underlying limitations in motor control, resulting greater dual-task costs and behavioral variability (Al-Yahya et al., 2011; McPhee et al., 2022). These measures may provide clinically informative markers of age-related functional and cognitive decline (Alzaid et al., 2022; Mustafovska et al., 2025).

Consistent with this view, we previously demonstrated that older adults exhibited greater motor costs and reaction-time variability during cognitive–motor dual-tasking than younger adults, while cognitive performance was relatively preserved (Deng et al., 2026). However, analyses based on performance averaged across an entire task condition treat dual-task interference as a relatively static phenomenon. They cannot reveal whether the initially elevated behavioral variability persists or changes as participants repeatedly engage with competing dual-task demands. Two bodies of evidence that capture dual-task performance at different timescales suggested the possibility of such within-session change. Acute dual-task studies primarily investigate the immediate performance costs produced by competing cognitive and motor demands (Bock, 2008; Bohle et al., 2019; Mustafovska et al., 2025). By contrast, dual-task practice and training studies demonstrate that these costs can be substantially reduced through repeated experience, with improvements observed in both motor and cognitive performance (Gallou-Guyot et al., 2020; Strobach, 2020; Zheng et al., 2021). Although examined separately from different timescales, these findings raised the possibility that dual-task performance may not be a static phenomenon but rather changes across multiple timescales, from initial interference to longer-term adaptation. However, it leaves an open question: whether meaningful adaptation can already emerge within the acute period itself. Such short-term changes would be obscured when performance is averaged across an entire task and characterized only in terms of its overall cost. Addressing this gap therefore requires examining how behavior and the dynamic network organization associated with it evolves over the course of a single dual-task session (Bassett et al., 2011; Mohseni et al., 2026).

Establishing whether behavior changes within a single dual-task session raises a corresponding neural question: how does network organization evolve alongside that behavior? Although previous studies have related neural fluctuations to performance (Kucyi et al., 2017; Seeburger et al., 2024), relatively little is known about how neural and behavioral dynamics unfold together during dual-task adaptation, particularly in aging. Performing two tasks concurrently requires coordination and reconfiguration across multiple functional systems as cognitive and motor demands interact and shift over time (Huang et al., 2017; Jiang et al., 2020; Worringer et al., 2019). This network organization may not remain constant throughout a dual-task session: configurations engaged during the initial response to interference may differ from those expressed later after participants gain experience with the competing demands (Mohseni et al., 2026; Strobach, 2020). These potential temporal phases refer not to different task conditions or to conventional pre- and post-training stages, but to changes unfolding within a single period of dual-task performance. Because static functional connectivity averages neural interactions across the entire session, it may obscure transient network reorganization associated with this process (Gbadeyan et al., 2022; Wang et al., 2024). Dynamic functional connectivity therefore provides a more appropriate framework for examining how network organization evolves in relation to behavioral adaptation (Bassett et al., 2011; Cohen, 2018; Finc et al., 2020).

Within this dynamic framework, dual-task performance can be conceptualized not as a single cost but as a potential adaptive trajectory. Initial task exposure may involve relatively high behavioral variability and competition between cognitive and motor demands. If performance becomes more stable with repeated experience, this change may be accompanied by reorganization of functional interactions across distributed brain networks. Studies of motor learning and working-memory training support this account by showing that behavioral improvement is accompanied by systematic network reconfiguration (Bassett et al., 2011; Finc et al., 2020). Moreover, older adults may adopt qualitatively different network organizations in response to increasing task demands (Iordan et al., 2021). However, it remains unknown whether comparable principles govern short-term cognitive–motor adaptation within a single dual-task session. The central question is therefore not only how large the initial dual-task cost is, but whether behavior and network organization change over the course of the session, and whether younger and older adults follow similar or distinct dynamic trajectories.

Answering these questions requires methods that preserve temporal information while capturing different properties of network organization. Dynamic independent component analysis (dyn-ICA) represents connectivity as continuously varying patterns of circuits expression, enabling the quantification of properties such as mean circuit strength and temporal variability (Alfonso Nieto-Castanon, 2020; Allen et al., 2014; Wang et al., 2024). Hidden Markov Models (HMM), by contrast, represent brain dynamics as transitions among recurring latent states, enabling the characterization of state occupancy and visitation patterns (Larsen et al., 2026; Stevner et al., 2019; Vidaurre et al., 2017). Task-based HMM studies have shown that these latent states exhibit distinct functional-connectivity configurations and that their occupancy, persistence, and switching behavior are associated with cognitive performance (Reddy et al., 2018; Taghia et al., 2018). These continuous and discrete approaches therefore provide related but non-equivalent descriptions of brain dynamics. Whereas dyn-ICA captures graded changes in the expression and variability of distributed connectivity patterns, HMMs identify recurring network configurations and characterize when and how frequently those configurations are visited. Previous studies have generally applied these perspectives separately (Hamid et al., 2025; Larsen et al., 2026; Wang et al., 2024), leaving unclear whether they identify overlapping or distinct features of behaviorally relevant network organization. Applying both methods to the same participants, task, and behavioral outcome enables a direct comparison within a common analytical framework. This combined approach allows us to determine whether continuous circuit properties and discrete state-visitation provide distinct or complementary correlates of dual-task adaptation, or point toward common principles of age-related dynamic network organization.

To address the limitations of condition-averaged analyses, the primary aim of the present study was to resolve the temporal trajectory of cognitive–motor dual-task performance within a single session. Specifically, we examined whether motor response-time variability remained stable or changed across repeated dual-task blocks and whether this trajectory differed between younger and older adults. We then investigated whether individual differences in within-session behavioral change were associated with dynamic functional-network organization, characterized through continuously varying circuit properties using dyn-ICA and recurring brain-state dynamics using HMM. Finally, we examined whether younger and older adults expressed distinct behaviorally relevant network profiles. By applying a common discovery and age-characterization framework to both approaches, we assessed whether continuous circuit expression and discrete state visitation identified overlapping or distinct features of the dynamic network organization associated with dual-task adaptation.

## 2 Methods

### 2.1 Participants

The participants included in the present study were drawn from the same cohort described in our previous study (Deng et al., 2026). The current work represents a secondary analysis of this dataset using a different analytical framework to investigates the brain functional dynamics and their relationship to dual-task behavioral adaptation. A total of 62 healthy, right-handed participants were recruited between 2023 and 2024 through university bulletin advertisements and local newspaper announcements. Participants were assigned to either a younger group (n = 20; 20–40 years) or an older group (n = 42; 50–80 years). The inclusion criteria were: (a) age within the predefined range for the respective group (20–40 years for younger adults and 50–80 years for older adults); (b) reported good physical and mental health, as assessed using the Short Form-12 Health Survey (SF-12), and the ability to walk independently without the use of a cane or walker; and (c) eligibility for MRI scanning (e.g., no metallic implants or claustrophobia). Exclusion criteria included a history of neurological, psychiatric, or movement disorders, current use of centrally acting medications, or any contraindication to MRI. The study was approved by the research ethics committee of the Carl von Ossietzky Universität Oldenburg and conducted in accordance with the declaration of Helsinki. All participants provided informed written consent. Table 1 summarizes the demographic characteristics and the final sample included in the present neuroimaging analyses. Detailed demographic characteristics, health status, and Test of Attentional Performance (TAP-M) behavioral assessments of this cohort have been reported previously (Deng et al., 2026). Only variables relevant to the present neuroimaging analyses are summarized here.

**Table 1.** Participant characteristics and MRI dataset used for dynamic functional connectivity and brain-behavior analyses.

| Characteristic | Older group<br>(n = 40) | Younger group<br>(n = 20) | p value |
| --- | --- | --- | --- |
| **Demographics** |  |  |  |
| Age (years) | 67.6 ± 7.1 | 28.0 ± 4.9 | <0.001 |
| Male, n (%) | 19 (47.5%) | 11 (55.0%) | 0.784 |
| Education (years) | 13.2 ± 3.1 | 14.6 ± 1.5 | 0.161 |
| **MRI data quality** |  |  |  |
| Mean framewise displacement (mm) | 0.16 ± 0.05 | 0.13 ± 0.02 | — |
| Subjects excluded due to excessive head motion | 2 | 0 | — |
| Subjects excluded due to missing behavioral data | 0 | 1 | — |
| Final sample for brain analyses | 40 | 20 | — |
| Final sample for brain-behavior analyses | 40 | 19 | — |

### 2.2 Experimental design

Task-based fMRI data comprised three task conditions: cognitive single-task, motor single-task, and cognitive–motor dual-task in a single functional run (Fig. 1). The order of the two single-task runs was counterbalanced across participants, whereas the dual-task run was always performed last. Each task began with a 10-s instruction period followed by eight experimental blocks, each lasting 25.8 s and separated by a 10-s rest period (Fig. 1). Every block consisted of 12 trials with a fixed trial duration of 2.15 s per trial, consisting of 1-s fixation period, a 600-ms stimulus presentation, and a 550-ms response window. This structure resulted in 96 trials across eight blocks, resulting in a total task duration of 296.4 s (≈ 4.94 min). The regular block structure enabled the quantification of block-wise characterization of motor behavior and corresponding changes in dynamic network organization across the dual-task session. A detailed description of the behavioral paradigm has been reported previously (Deng et al., 2026). The key features relevant to the present analyses are summarized below.

**Figure 1.**
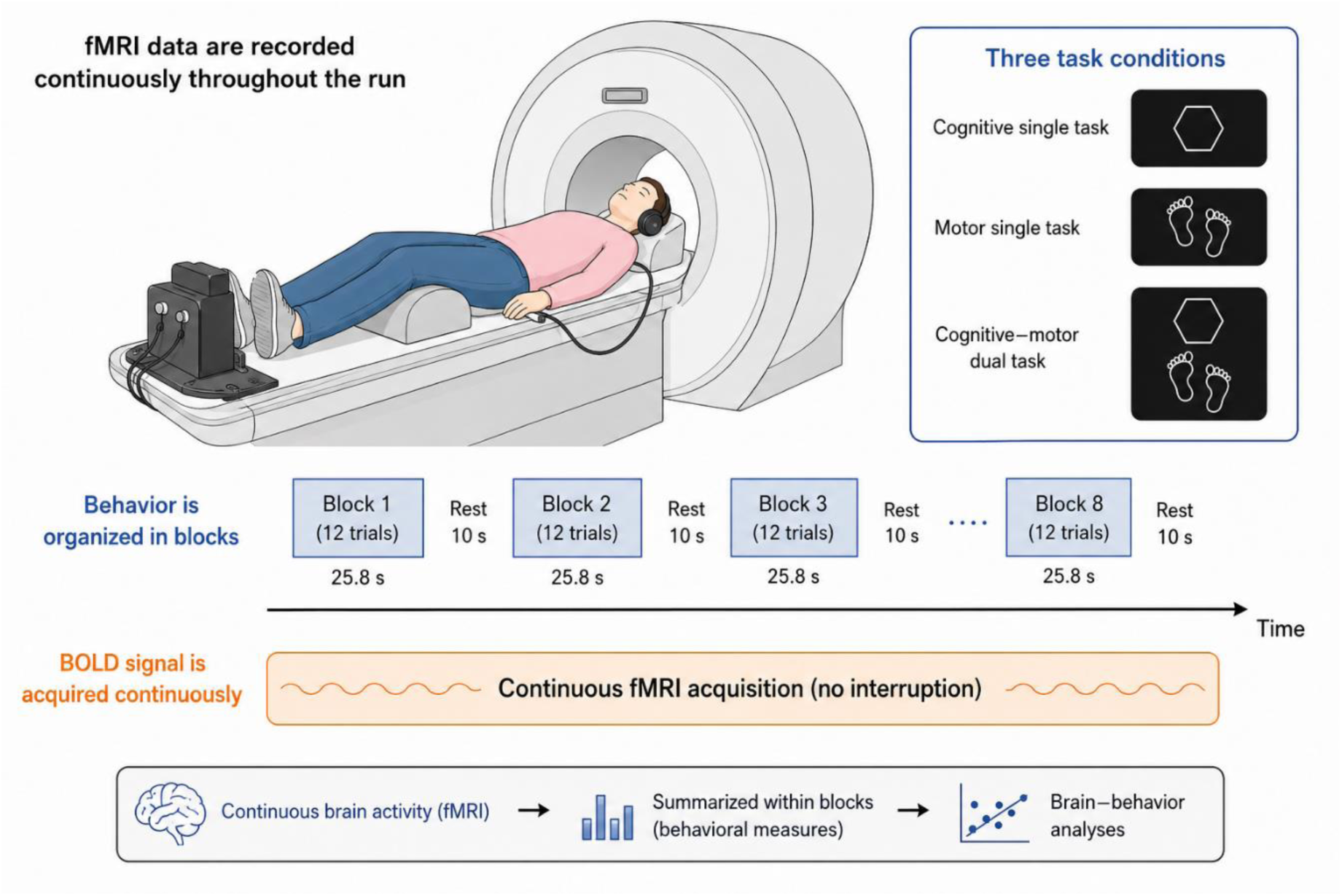
Experimental paradigm and block-wise analytical framework. Task-based fMRI data were acquired continuously throughout a single functional run comprising three conditions: cognitive single-task, motor single-task, and cognitive–motor dual-task. Each condition contained eight 25.8-s experimental blocks separated by 10-s rest periods, with12 trials of 2.15 s in each block. Data from the whole functional run comprising the three task conditions was used for the estimation of the common dynamic network models. For the primary analyses of within-session dual-task adaptation, motor behavioral performance and dynamic network metrics were summarized within each dual-task block and examined using brain–behavior associations. Motor reaction-time variability during the single-motor condition served as a reference for interpreting dual-task performance.

In the cognitive single task, participants performed a Go/NoGo paradigm adapted from the Human Connectome Project in Aging (Bookheimer et al., 2019). Six geometric shapes served as Go stimuli requiring a right-index-finger button press, whereas two shapes (square and circle) served as NoGo stimuli requiring response inhibition. Go and NoGo trials were presented in a randomized order with a 75:25 ratio. In the motor single task, participants performed rhythmic alternating left- and right-foot movements using an MRI-compatible pedaling device in response to a foot cue presented on the lower half of the screen. The stepping frequency matched the cognitive task timing, with one step performed every 2.15 s. In the **cognitive–motor dual task**, the cognitive and motor stimuli were presented simultaneously in the upper and lower halves of the screen, respectively. During Go trials, participants performed both a button press and a foot movement; during NoGo trials, they withheld the button press while continuing the rhythmic stepping movement. The identical trial timing and block structure across conditions allowed motor variability during the single-motor condition to provide a task-matched reference for interpreting performance during the dual-task run. All tasks were presented and cognitive and motor responses were recorded, using Presentation software (version 22.1; Neurobehavioral Systems, Berkeley, CA, USA).

Task-based fMRI data from the whole functional run was used to estimate a common dynamic network space comprising the dyn-ICA circuits and HMM states. The primary behavioral outcomes were dual-task motor reaction time and motor reaction-time variability across the eight dual-task blocks. Motor performance during the single-motor condition was used only as a reference for interpreting dual-task performance, whereas cognitive performance during the single-cognitive condition was not included in the present analyses.

### 2.3 fMRI data acquisition

Task-based fMRI data were acquired using a 3-T Siemens MAGNETOM Prisma scanner (Siemens Healthcare, Erlangen, Germany) with a 64-channel head coil. Participants were instructed to lie comfortably on the scanner bed, and the pedal device was individually adjusted to permit a natural range of leg extension and comfortable foot movements. To minimize task-related head motion, participants were instructed to keep their heads still during pedaling. Soft foam pads were placed beneath the head and knees for stabilization. In addition, a strip of medical tap (Leukosilk) was placed across the forehead and attached to the coil borders to provide tactile feedback about head movement.

Before the main task run, participants completed an approximately 4.5-min practice session covering the single- and dual-task conditions, followed by a brief performance check and feedback. The cognitive single-task, motor single-task, and cognitive–motor dual-task conditions were then performed within a single continuous functional run (Fig. 1). The order of the two single-task conditions was counterbalanced across participants, whereas the dual-task condition was always presented last. Functional images were then acquired continuously throughout the complete task using a T2*-weighted multiband echo planar imaging (EPI) sequence with BOLD contrast. Acquisition parameters were as follows: repetition time (TR) = 850 ms, echo time (TE) = 30 ms, flip angle = 62°, slice thickness = 2.5 mm, field of view = 192×192 mm, acquisition matrix = 76×76, voxel size = 2.5×2.5×2.5 mm³, 48 slices, multiband acceleration factor = 4. The functional run lasted 15 min 26 s and comprised 1090 volumes. Following the functional run, a high-resolution T1-weighted structural image was acquired using the following parameters: TR = 2000 ms, TE = 2.07 ms, flip angle = 9°, voxel size = 0.75×0.75×0.75 mm³, GRAPPA acceleration factor = 2, field of view = 240×240 mm, 224 sagittal slices, and acquisition time = 6 min 16 s.

### 2.4 fMRI data preprocessing

Functional MRI data were preprocessed and denoised using the CONN toolbox (RRID: SCR_009550, version 22.v2407; Nieto-Castanon, 2020; Whitfield-Gabrieli & Nieto-Castanon, 2012) implemented in MATLAB R2023a together with SPM12 (RRID: SCR_007037, version 12.7771). Preprocessing followed the default CONN pipeline and included functional realignment and unwarping, slice-timing correction, artifact detection, tissue segmentation, normalization to the MNI152 template (2-mm isotropic resolution), and spatial smoothing using an 8-mm full-width-at-half-maximum (FWHM) Gaussian kernel.

Potential motion-related outlier volumes were identified using the Artifact Detection Tools (ART) with thresholds of framewise displacement > 0.9 mm or global BOLD signal changes > 5 standard deviations. Functional data were subsequently denoised using linear regression of nuisance signals, including five anatomical CompCor (aCompCor) components each from white matter and cerebrospinal fluid, 12 head-motion regressors (six rigid-body motion parameters and their temporal derivatives), task effects and their first-order derivatives (10 regressors), linear trends, and one regressor for each identified outlier volume (Power et al., 2014). On average, 22 of 1,090 volumes per participant were censored. Task-related regressors were included to remove task-evoked activation, thereby ensuring that subsequent functional connectivity estimates reflected condition-dependent interactions rather than coactivation effects. Finally, the residual BOLD time series were band-pass filtered between 0.008 and 0.09 Hz, resulting in effective degrees of freedom ranging from 94.5 to 147.2 (mean = 143.1). The resulting denoised BOLD time series served as the input for subsequent dynamic functional connectivity analyses using both continuous (dynamic ICA) and discrete-state (Hidden Markov Model) approaches. All fMRI data preprocessing and dynamic functional connectivity modeling were performed on the high-performance computing cluster at Carl von Ossietzky Universität Oldenburg.

### 2.5 Dynamic functional connectivity

To investigate how dynamic functional brain organization supports behavioral adaptation during dual-task performance, we applied two complementary approaches. Dynamic independent component analysis (dyn-ICA) was used to characterize continuously varying patterns of functional connectivity (Nieto-Castanon, 2020; Hamid et al., 2025; Wang et al., 2024), whereas Hidden Markov Modeling (HMM) was used to identify recurring latent brain states and characterize their temporal organization (Larsen et al., 2026; Stevner et al., 2019; Vidaurre et al., 2017).

To enable direct methodological comparison, dyn-ICA and HMM were evaluated using a common discovery and age-characterization framework (Table 2). First, each method was estimated independently at the group level using data from all participants and all three task conditions within the single functional run. Dyn-ICA identified dynamic connectivity circuits, whereas HMM identified recurring latent states. Second, the group-level representations were mapped back to individual participants, yielding subject-specific circuit time courses for dyn-ICA and state-probability time courses for HMM. Third, two subject-level metrics were derived for each circuit or state and tested for associations with dual-task adaptation scores using Pearson correlations. Multiple comparisons were controlled separately within each methodological framework using the Benjamini–Hochberg false discovery rate (FDR) procedure across 20 discovery tests. Finally, age-group differences were examined across all ten circuits and all ten states, rather than only the behaviorally relevant features. Applying the same analytical framework to both methods enabled direct comparison of continuous circuit expression and discrete state dynamics while preserving their distinct mathematical representations. This comparison allowed us to assess whether the two approaches identified overlapping or distinct neural correlates of within-session dual-task adaptation and whether they converged on common principles of age-related dynamic network organization.

**Table 2.**
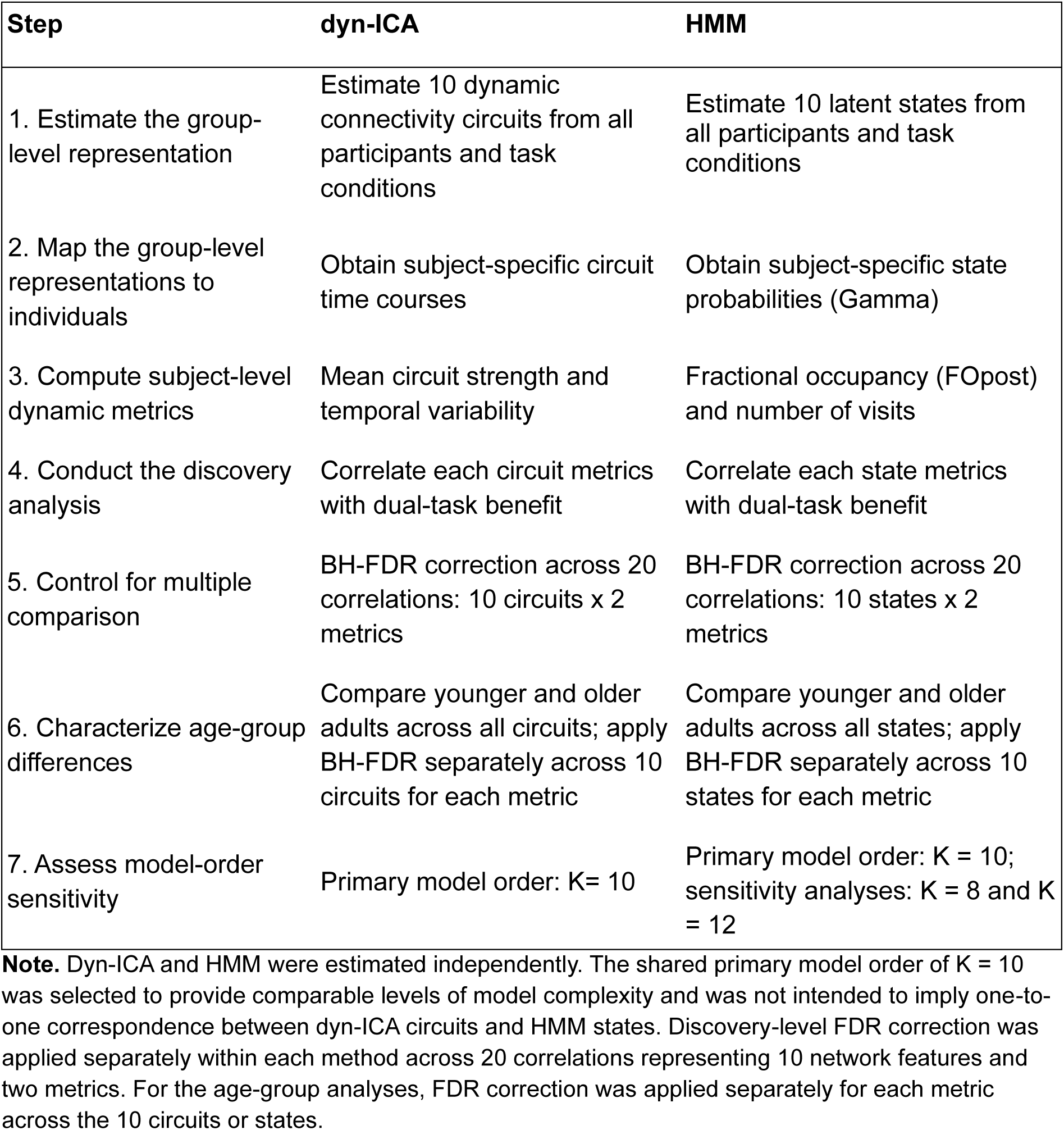
Common analytical framework for continuous dyn-ICA and discrete HMM analyses.

| Step | dyn-ICA | HMM |
| --- | --- | --- |
| 1. Estimate the group-level representation | Estimate 10 dynamic connectivity circuits from all participants and task conditions | Estimate 10 latent states from all participants and task conditions |
| 2. Map the group-level representations to individuals | Obtain subject-specific circuit time courses | Obtain subject-specific state probabilities (Gamma) |
| 3. Compute subject-level dynamic metrics | Mean circuit strength and temporal variability | Fractional occupancy (FOpost) and number of visits |
| 4. Conduct the discovery analysis | Correlate each circuit metrics with dual-task benefit | Correlate each state metrics with dual-task benefit |
| 5. Control for multiple comparison | BH-FDR correction across 20 correlations: 10 circuits x 2 metrics | BH-FDR correction across 20 correlations: 10 states x 2 metrics |
| 6. Characterize age-group differences | Compare younger and older adults across all circuits; apply BH-FDR separately across 10 circuits for each metric | Compare younger and older adults across all states; apply BH-FDR separately across 10 states for each metric |
| 7. Assess model-order sensitivity | Primary model order: K= 10 | Primary model order: K = 10; sensitivity analyses: K = 8 and K = 12 |

**Note.** Dyn-ICA and HMM were estimated independently. The shared primary model order of K = 10 was selected to provide comparable levels of model complexity and was not intended to imply one-to-one correspondence between dyn-ICA circuits and HMM states. Discovery-level FDR correction was applied separately within each method across 20 correlations representing 10 network features and two metrics. For the age-group analyses, FDR correction was applied separately for each metric across the 10 circuits or states.

#### Model-order selection and sensitivity analysis

For the primary analyses, the model order was set to K=10 for both dyn-ICA and HMM. This intermediate model order was selected a priori to provide sufficient differentiation among dynamic network configurations while limiting fragmentation into potentially unstable or difficult-to-interpret components. Importantly, K was specified before conducting the brain–behavior discovery analyses and was not selected on the basis of the strength or statistical significance of the resulting behavioral associations. Using the same primary model order also facilitated direct comparison between the continuous and discrete representations, although the two methods were estimated independently and were not assumed to produce one-to-one corresponding components. To assess whether the principal findings depended on the selected model order, the sensitivity analyses were conducted using K=8 and K=12. We evaluated whether the alternative solutions reproduced the principal behavioral associations observed in the K=10 solution. These sensitivity analyses yielded comparable main patterns, indicating that the central conclusions were not specific to a single choice of K (Supplementary Material: Figures S1–S4).

#### 2.5.1 Continuous network dynamics

To quantify continuous changes in large-scale functional connectivity during task performance, we applied dynamic independent component analysis (dyn-ICA) as implemented in the CONN toolbox, to the preprocessed fMRI data (Nieto-Castanon, 2020). Dyn-ICA combines generalized psychophysiological interaction (gPPI) modeling with ICA-based decomposition to identify groups of functional connections that exhibit similar temporal modulation throughout the functional run.

Two distinct uses of ICA should be distinguished. First, the functional network space was defined using 32 regions of interest (ROIs) from the CONN network atlas (Nieto-Castanon, 2020). These ROIs were derived independently from ICA of Human Connectome Project data from 497 participants and included the default-mode (4 ROIs), sensorimotor (3), visual (4), salience/cingulo-opercular (7), dorsal-attention (4), frontoparietal/central-executive (4), language (4), and cerebellar (2) networks. Thus, defining the ROI space did not involve performing ICA on the present sample. Denoised BOLD time series extracted from these 32 ROIs served as the input to the dynamic connectivity analysis. Full ROI definitions are provided in Supplementary Table S1.

Second, dyn-ICA was applied to the ROI-to-ROI connectivity data from the present participants. Temporal modulation of connectivity between each ROI pair was estimated within a gPPI framework (Friston et al., 1997). CONN’s iterative dual-regression procedure was then used to estimate temporal modulation factors representing continuous changes in connectivity. At the group level, ICA decomposed these modulation patterns into statistically independent dynamic connectivity circuits. Each circuit was defined by a characteristic pattern of connections among the 32 ROIs and an associated time course of circuit describing its expression throughout the functional run.

Following the default CONN implementation, temporal smoothing was performed using a 30-s full-width-at-half-maximum (FWHM) Hanning kernel. The resulting dynamic circuits represent recurring patterns of functional connectivity, whereas their associated temporal modulation factors describe the continuous expression of each circuit over time. For each participant, dyn-ICA generated a continuous time course for each of the 10 dyn-ICA circuits across the entire fMRI run. These subject-specific time courses were subsequently used to derive summary measures of mean circuit strength and temporal variability for the brain–behavior analyses (Table 2).

#### 2.5.2 Discrete network dynamics

To characterize transitions among recurring functional brain configurations, we applied a Gaussian Linear Hidden Markov Model (GLHMM) following the protocol described by Larsen et al., (2026). The denoised BOLD time series from the same 32 ROIs used in the dyn-ICA analysis served as input to the HMM. For each participant, the data were arranged as a time-points × ROI-features matrix and then concatenated across participants to form a group-level dataset. A single HMM was therefore estimated across the complete sample. This procedure defined a common latent state space, allowing state expression to be compared directly across participants, conditions, and age groups. Before model estimation, the data were standardized according to the GLHMM protocol (Larsen et al., 2026).

For the primary analysis, the model order was specified as K=10 latent states before conducting the brain–behavior discovery analysis. This intermediate model order was selected to provide differentiation among recurring brain configurations while limiting excessive fragmentation into potentially unstable or difficult-to-interpret states. Its correspondence with the 10-circuit dyn-ICA solution additionally facilitated comparison between the two approaches but did not imply a one-to-one mapping between circuits and states. Model parameters were estimated using variational inference implemented in the GLHMM toolbox. The fitted model yielded posterior state-probability time courses (Gamma), representing the probability that each latent state was occupied at every time point. Robustness to model order was evaluated using sensitive analyses with K=8 and K=12, as described in Section 2.5, Model-order selection and sensitivity analysis.

Analogous to the mapping-back procedure used for dyn-ICA, the group-level HMM representation was mapped back to each participant and experimental condition according to the original time-series boundaries. This enabled participant- and condition-specific HMM metrics to be calculated while preserving a common set of state definitions across the entire sample.

Three standard metrics were extracted for each state: posterior fractional occupancy (FOpost), defined as the mean posterior probability of occupying that state; mean lifetime, defined as the average duration of consecutive state visits in seconds; and number of visits, defined as the frequency with which a state was entered. Because fractional occupancy and mean lifetime both quantify state persistence and are typically highly correlated, the primary discovery analysis was restricted to two complementary metrics: fractional occupancy (FOpost) and number of visits. Mean lifetime was retained for secondary descriptive characterization of behaviorally relevant states but was excluded from the discovery-stage multiple-comparison correction to reduce redundancy.

#### 2.5.3 Discovery of behaviorally relevant dynamic network features

To characterize within-session behavioral change, motor RT variability was calculated separately within each of the eight dual-task blocks as the standard deviation of motor RT across the 12 trials in that block. Behavioral adaptation was quantified using a signed dual-task adaptation score, defined as the percentage reduction in RT variability relative to the initial dual-task block:

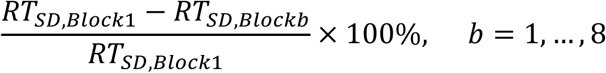

Here, *RT_SD,Block1_* represents motor RT variability during Block 1, and *RT_SD,Blockb_* represents variability during the corresponding block. Positive scores therefore indicate reduced RT variability and greater motor stabilization relative to Block 1, hereafter referred to as dual-task benefit; zero indicates no change; and negative scores indicate increased variability. By definition, dual-task adaptation at Block 1 was zero. The directional formulation follows the relative-change principles applied to dual-task effects by Kelly et al. (2010), whereby change is expressed relative to a reference value and oriented so that positive scores indicate better performance. Unlike a conventional dual-task effect, which compares average dual-task performance with a single-task reference, the present measure compares repeated blocks within the same dual-task condition. It therefore quantifies within-session adaptation relative to each participant’s initial dual-task performance, allowing behavioral change to be traced across the entire dual-task session.

For the subsequent brain–behavior analyses, dual-task adaptation score at the final block (Block 8) was used as the summary measure of overall within-session adaptation, representing the relative change in motor RT variability from the beginning to the end of the dual-task condition. All dyn-ICA circuits and HMM states were systematically screened for associations with individual differences in this endpoint measure. For dyn-ICA, subject-level metrics describing the mean circuit strength and temporal variability were computed for each of the ten dynamic circuits. For HMM, fractional occupancy (FOpost) and number of visits were calculated separately for each state within each dual-task block. For each dynamic feature, Pearson correlation coefficients were calculated between the subject-level metric and dual-task adaptation score. If the score is positive, we refer it as dual-task benefit. This resulted in 20 statistical tests for dyn-ICA (10 circuits × 2 metrics) and 20 tests for HMM (10 states × 2 metrics). Multiple comparisons were controlled separately for each method using the Benjamini–Hochberg false discovery rate (FDR) procedure. Dynamic circuits and brain states were ranked according to their FDR-adjusted *q* values.

#### 2.5.4 Characterization and age-related expression of behaviorally relevant network feature

Dynamic circuits and HMM states identified in the discovery analysis were further characterized in relation to behavioral adaptation and age. For the focused visualization of each behaviorally relevant feature, we presented the network metric through which that feature had been identified during discovery and its Pearson correlation with dual-task benefit. These plots illustrated the strength and direction of the previously identified brain–behavior relationships and were not treated as independent confirmatory tests.

To understand the broader age-related expression in both dynamic circuits and HMM states we examined their age-group difference at two levels. First, the behaviorally relevant features were compared between younger and older adults to determine whether the neural signatures associated with dual-task adaptation differed by age. Second, the analysis was extended across all ten dyn-ICA circuits and all ten HMM states to establish whether age differences in the behaviorally relevant features were selective or formed part of a broader network-wide pattern. Age-group differences were assessed using Welch’s two-sample t-tests, which accommodate unequal group sizes and variances. Multiple comparisons were controlled using the Benjamini–Hochberg false discovery rate procedure, applied separately across the ten dyn-ICA circuits for mean strength and temporal variability and across the ten HMM states for fractional occupancy (FOpost) and number of visits. Group means, standard deviations, standard errors of the mean, older-minus-younger mean differences, and Cohen’s *d* effect sizes were calculated for each comparison.

Age-group comparisons of brain measures included all 60 participants (20 younger and 40 older adults), including the participant for whom behavioral data were unavailable. Brain–behavior analyses included 59 participants with both imaging and behavioral data (Table 1). Together, these analyses allow us to test whether successful dual-task adaptation is supported by specific dynamic network configurations and whether age-related differences in behavioral trajectories reflect distinct network-level strategies for achieving behavioral stability.

## 3. Results

### 3.1 Dynamic functional connectivity and behavioral adaptation

Figure 2 provides a representative single-subject illustration of the temporal relationship between dynamic functional connectivity (FC) and motor performance during the dual-task condition, before presenting the group-level analyses in Figures 3–10. The dual-task period comprised eight consecutive task blocks (shaded regions), each containing 12 trials, resulting in a total of 96 trials (Figure 2A). The example highlights the moment-to-moment fluctuations in dynamic FC alongside trial-wise motor stepping RT across the dual-task session. The corresponding block-wise motor RT variability for the same participant is shown in Figure 2B. RT variability was highest during the first dual-task block and progressively decreased across Blocks 2–7, reaching its lowest value in Block 7 before exhibiting a slight increase in the final block (Block 8). This example illustrates the concurrent evolution of dynamic FC and motor behavior across the entire dual-task session. Group-level analyses are presented next to determine whether this adaptation pattern is consistently observed across participants.

**Figure 2.**
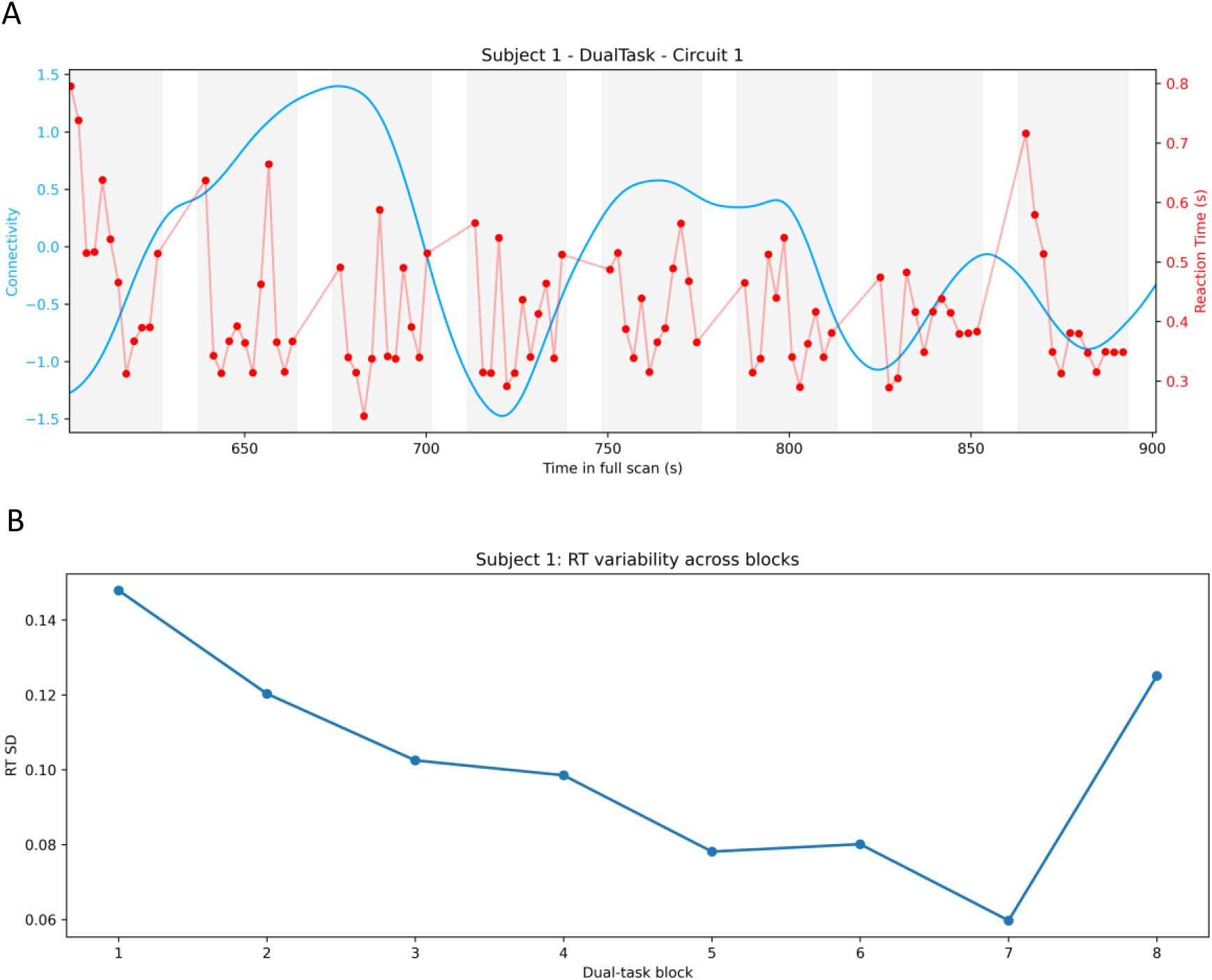
Single-subject example of dynamic functional connectivity and trial-wise motor reaction time variability during dual-task performance. (A) Time course of dynamic functional connectivity (blue line; Circuit 1, subject 1) and trial-wise motor stepping reaction times (red dots and line) during the dual-task condition. The shaded regions indicate the eight consecutive task blocks (12 trials per block). (B) Block-wise motor reaction time variability (standard deviation of reaction times) for the same subject.

**Figure 3.**
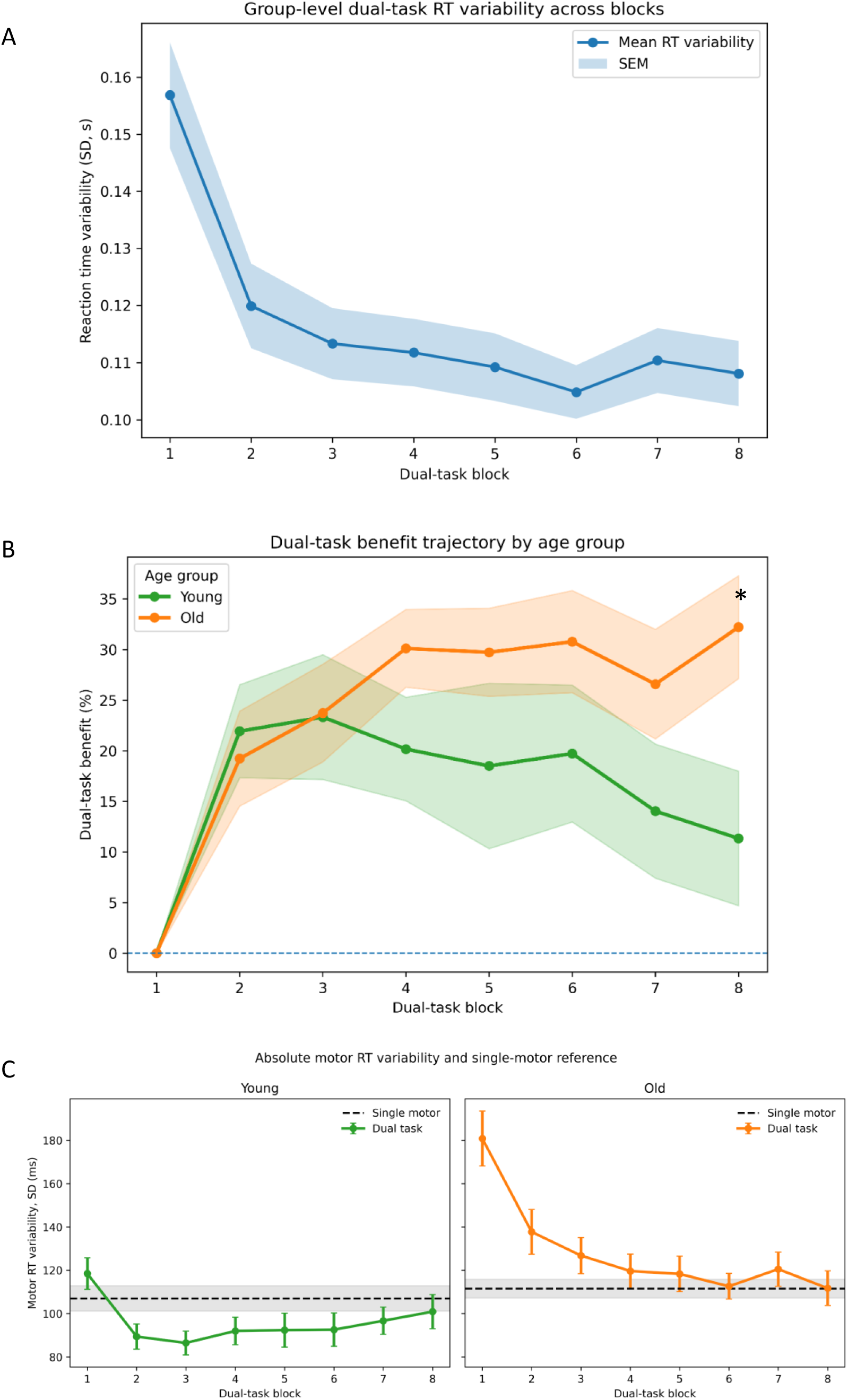
Group-level evolution of dual-task reaction time variability across dual-task blocks. (A) Group mean motor reaction time variability (standard deviation of reaction times) across the eight dual-task blocks is shown, with shaded areas indicating the standard error of the mean (SEM). (B) Adaptive dual-task benefit trajectories for older (orange line) and younger (green line) groups. Motor reaction time variability was initially high but progressively decreased with repeated dual-task exposure, indicating adaptive stabilization of performance. Older adults exhibited a larger adaptive dual-task benefit (∼30%) than younger adults (∼10–20%). Asterisk marks a significant between-group difference at the final block (p = 0.017). (C) absolute RT variability in young (left panel) and old group (right panel). The dashed lines indicate the single-motor reference for each group.

**Figure 4.**
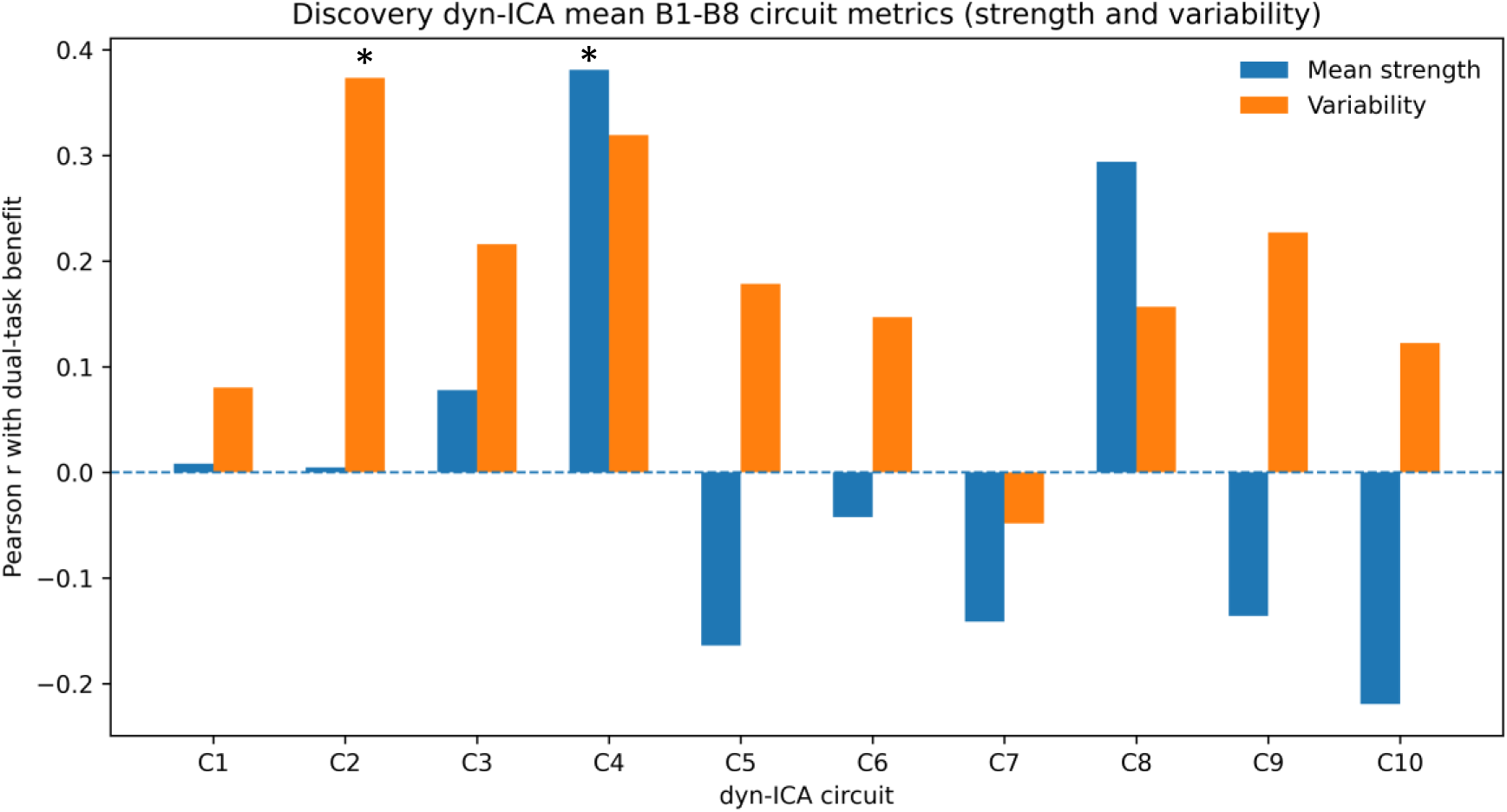
Discovery of behaviorally relevant dyn-ICA circuits. Bar plots show Pearson correlations coefficients (r) between dual-task benefit and two dyn-ICA circuits metrics (circuit strength and temporal variability) across the ten circuits. Positive correlations indicate that greater circuit expression was associated with larger improvements in dual-task performance, while negative correlations predict worse performance. Circuits surviving FDR correction (*q* < 0.05, 20 tests) are marked with asterisks: Circuit 4 mean strength (*r* = 0.38, *q* = 0.036), and Circuit 2 temporal variability (*r* = 0.37, *q* = 0.036). Significance: * *q* < 0.05.

**Figure 5.**
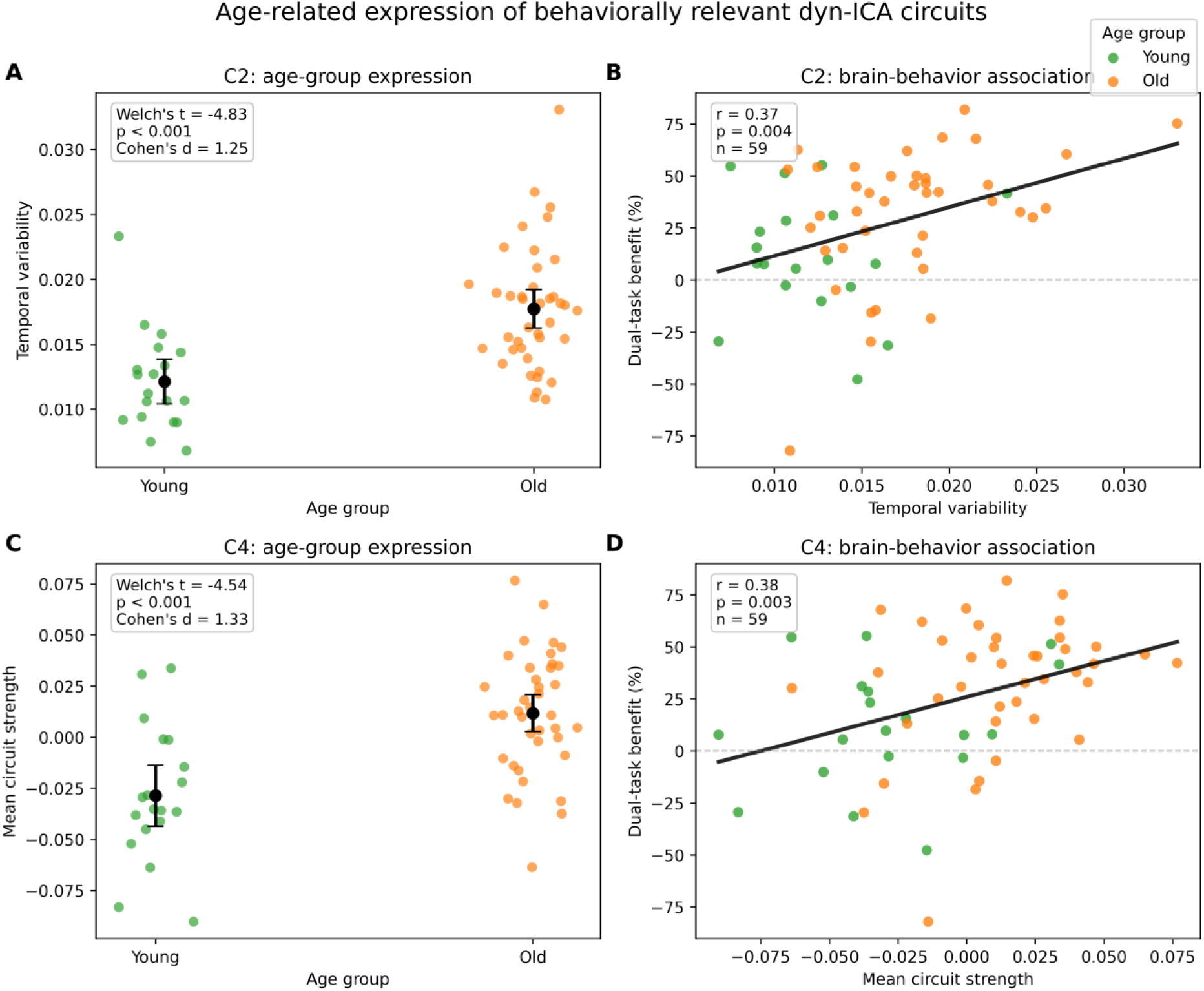
Age-group differences in the two behaviorally relevant dyn-ICA features. (A) Age-group difference in Circuit 2 temporal variability. (B) Association between Circuit 2 temporal variability and dual-task benefit across all participants. (C) Age-group difference in Circuit 4 mean strength. (D) Association between Circuit 4 mean strength and dual-task benefit across all participants.

**Figure 6.**
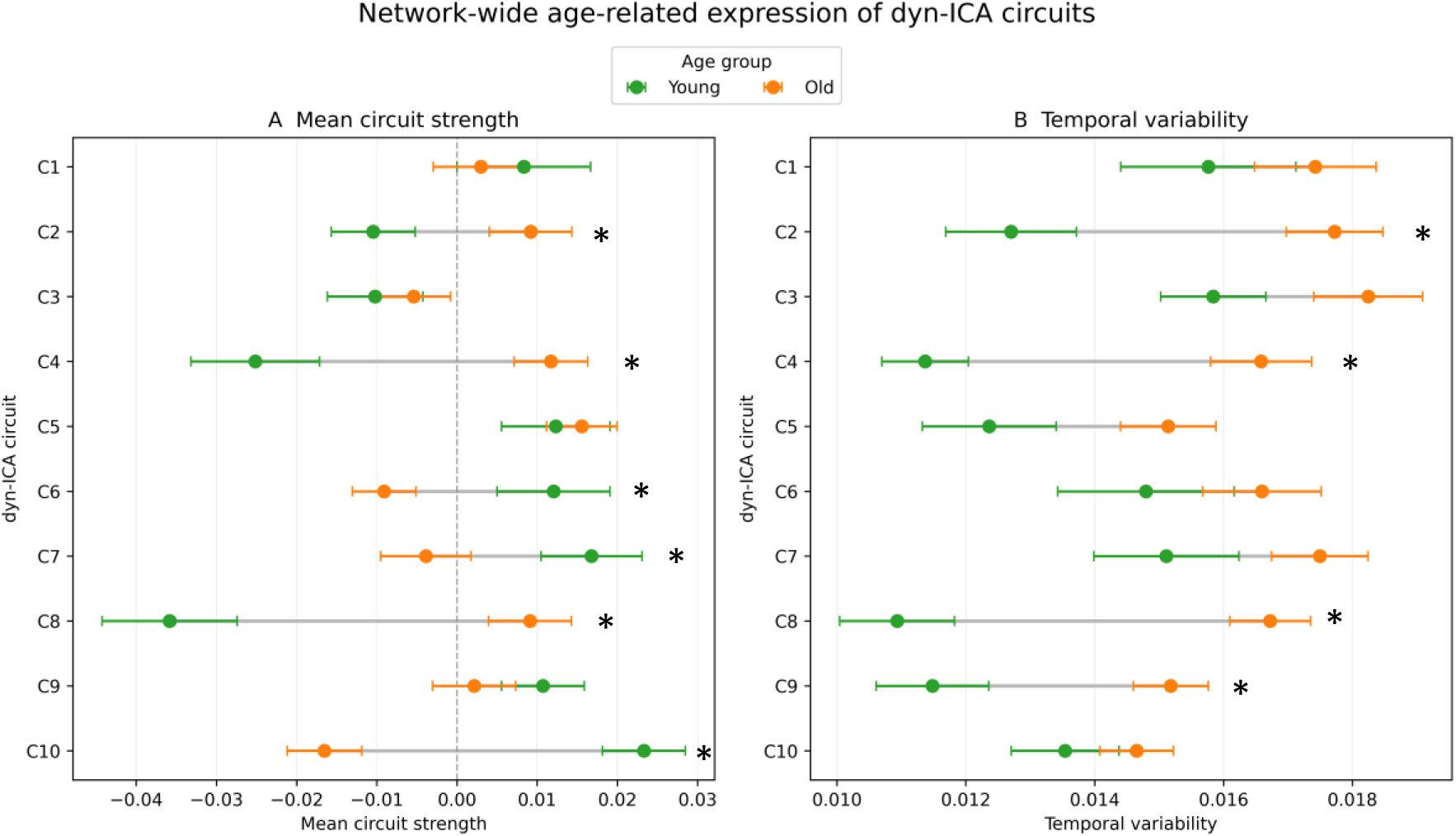
Network-wide age-group differences in dyn-ICA circuit expression. Bar plots show the mean ± SEM for younger and older adults across all ten dyn-ICA circuits for (A) mean circuit strength and (B) temporal variability. * marks a significant between-group difference after FDR correction.

**Figure 7.**
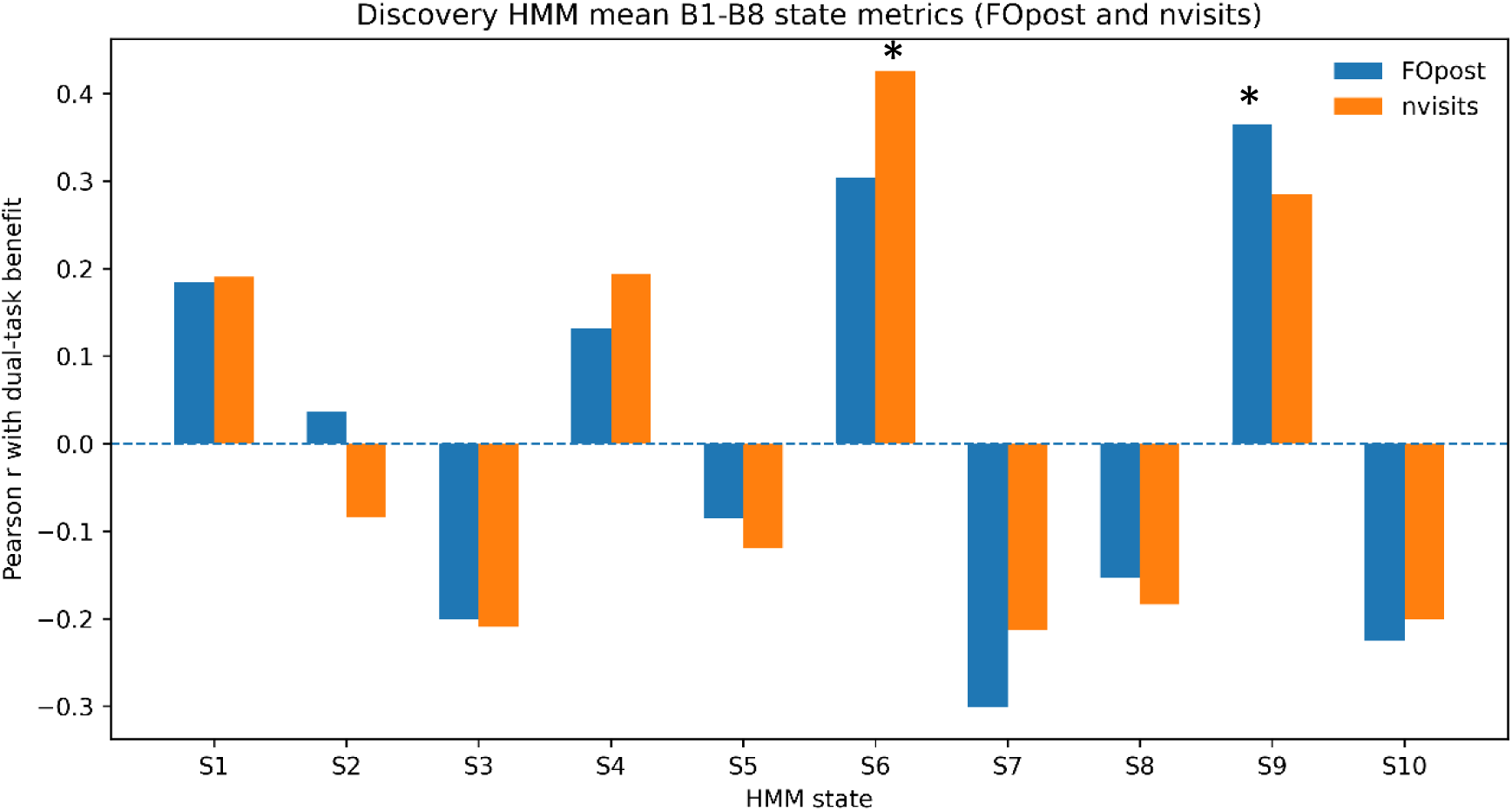
Discovery of behaviorally relevant HMM states. Bar plots show Pearson correlations coefficients (r) between dual-task benefit and two HMM state metrics (fractional occupancy, and number of visits) across the ten states. Positive correlations indicate that greater state engagement was associated with larger improvements in dual-task benefits, while negative correlations predict worse performance. States surviving FDR correction (*q* < 0.05, 20 tests) are marked with asterisks: State 6 number of visits (*r* = 0.43, *q* = 0.015), and State 9 fractional occupancy (*r* = 0.37, *q* = 0.044). Significance: * *q* < 0.05.

**Figure 8.**
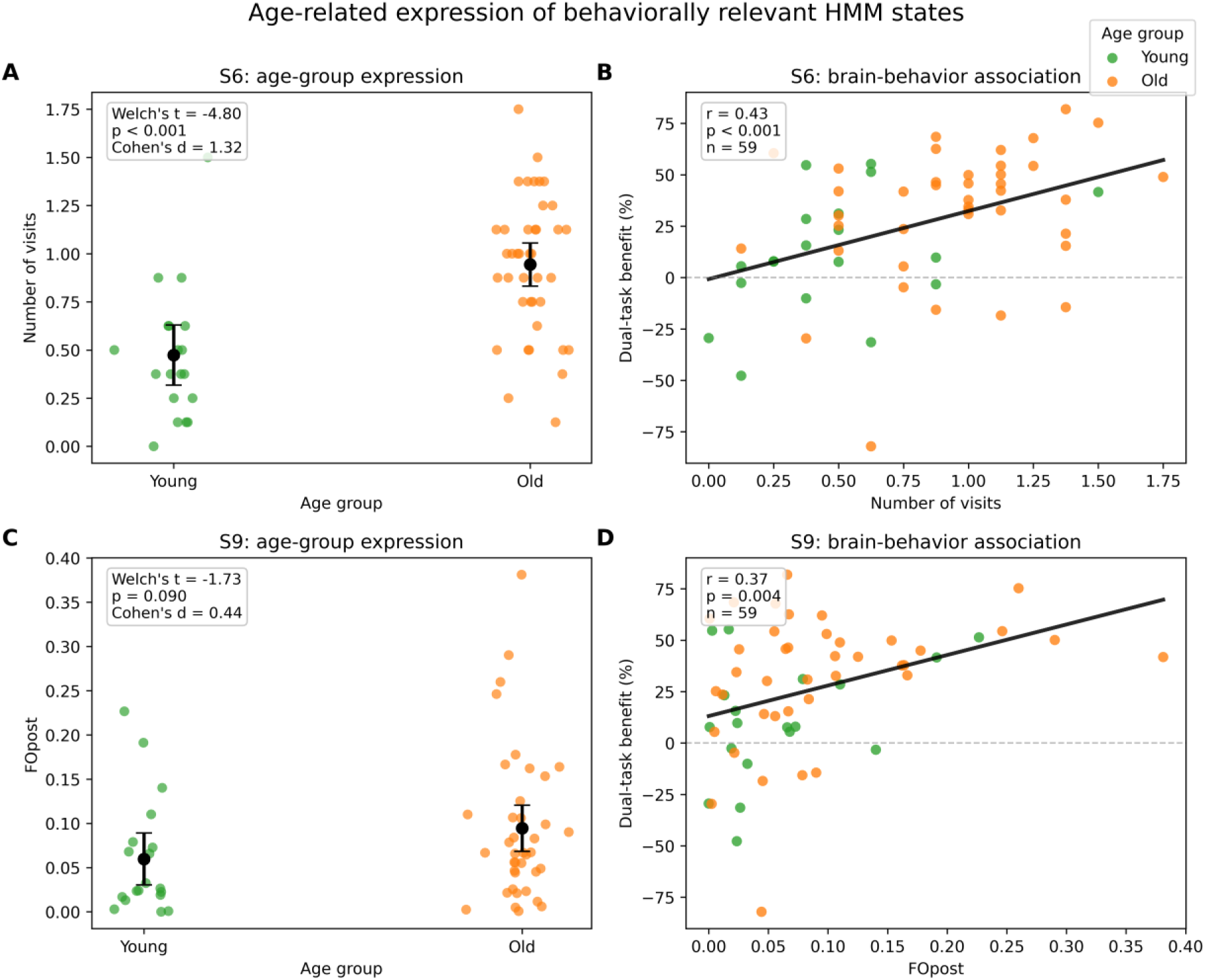
Age-group differences in the two behaviorally relevant HMM features. (A) Age-group difference in State 6 number of visits (B) Association between State 6 number of visits and dual-task benefit across all participants. (C) Age-group difference in State 9 fractional occupancy. (D) Association between State 9 fractional occupancy and dual-task benefit across all participants.

**Figure 9.**
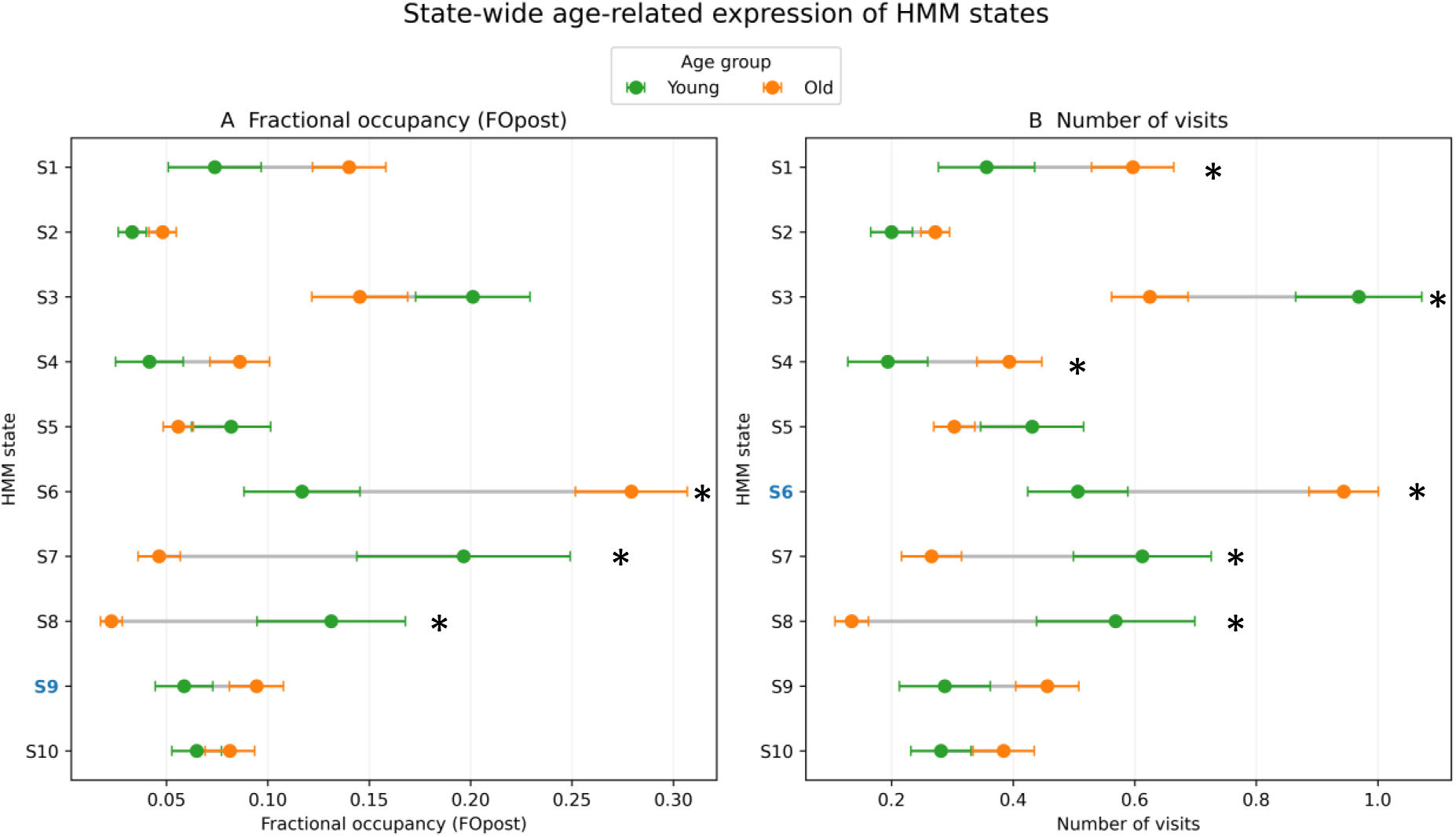
Age-group differences in all ten HMM state expression. Bar plots show the mean ± SEM for younger and older adults across all ten HMM states for (A) fractional occupancy and (B) number of visits. State and metric with age-group difference surviving FDR correction are marked with asterisks. The behaviorally relevant features identified in Figure 7—State 6 number of visits and State 9 fractional occupancy —are highlighted with blue color of labels.

**Figure 10.**
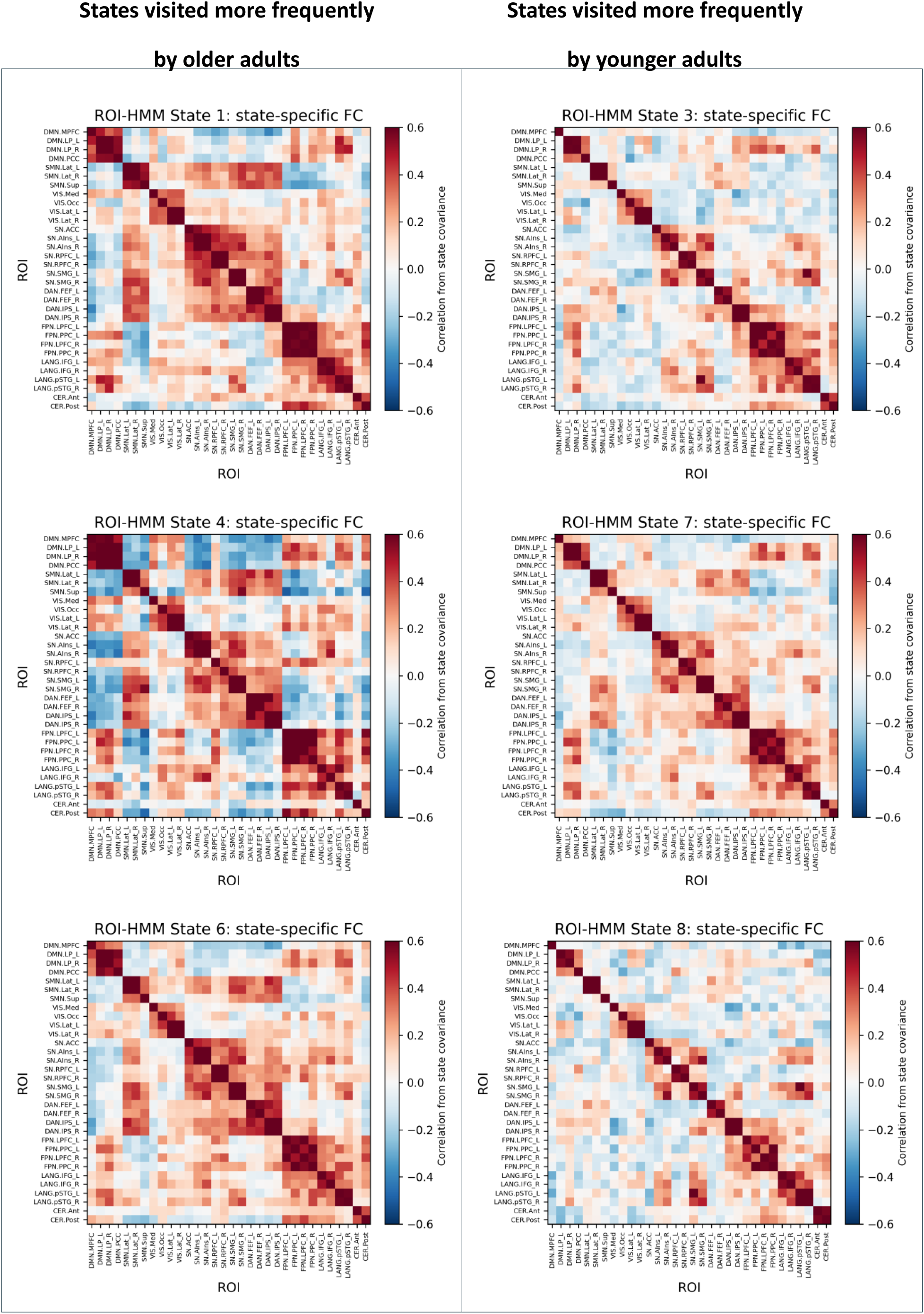
Connectivity configurations of HMM states showing age-related differences in visitation. State-specific functional connectivity matrices are shown for the six HMM states whose number of visits differed significantly between younger and older adults after Benjamini–Hochberg FDR correction across all ten states. States visited more frequently by older adults are shown in the left column (States 1, 4, and 6), whereas states visited more frequently by younger adults are shown in the right column (States 3, 7, and 8). Matrices display the state-specific functional connectivity configurations estimated from the HMM fitted jointly across all participants. Red and blue values indicate positive and negative state-specific connectivity, respectively. Column assignment therefore reflects age-group differences in state visitation, not connectivity matrices estimated separately within each age group.

The group-level evolution of dual-task RT variability is shown in Figure 3A. Consistent with the single-subject example, RT variability was highest during the first dual-task block and progressively decreased across subsequent blocks, reaching a minimum at Block 6. A slight increase was observed in Block 7, with variability remained relatively stable in Block 8. This temporal profile suggests a dynamic adaptation process characterized by high initial motor variability and gradual stabilization of performance over repeated dual-task exposure.

Dual-task adaptation score was calculated separately for each participant and each block as the percentage reduction in motor RT variability relative to the initial dual-task block (Block 1). The larger positive scores indicate greater improvement, and hence refer to as dual-task benefit (see Methods, Section 2.5.3). These individual benefit values were then summarized by age group to characterize the trajectories of older and younger adults (Figure 3B). Both groups exhibited comparable improvements during the early phase of adaptation (Blocks 2–3), after which their trajectories diverged. From Block 4 onward, older adults generally maintained a greater relative reduction in RT variability, with dual-task benefit reaching approximately 30% by the final block. In contrast, benefit among younger adults decreased to approximately 10–20%, reaching 11.34% at Block 8. The between-group difference was statistically significant at the final block (p = 0.017).

To contextualize this apparent age-related divergence in relative dual-task benefit, we compared absolute motor variability during each dual-task block with the corresponding age group’s single-motor references. Younger adults began only slightly above their single-motor reference, showed a rapidly reduction by Block 2, and maintained lower variability than their reference throughout Blocks 2–8. In contrast, older adults began with markedly elevated variability during Block 1, followed by a pronounced early reduction at Block 2 and progressively approached their single-motor reference, which they essentially reached by Blocks 6 and 8. These distinctive trajectories help explain the larger percentage benefit observed in older adults: they experienced much greater initial instability and therefore had more room for improvement. Their approximately 32% benefit reflects a substantial recovery toward their own single-motor baseline. Conversely, younger adults started closer to single-motor baseline and quickly became even less variable than during the single-motor condition. Thus, by the end of session, older adults had largely returned to their single-motor stability, whereas younger adults had maintained sub-reference motor variability across most of the dual-task condition.

### 3.2 Continuous network dynamics (dyn-ICA)

To identify dynamic network features associated with dual-task performance, we correlated two metrics from each analytical framework with dual-task benefit, defined as the relative positive reduction in motor RT variability from the beginning (Block 1) to the end of the dual-task condition (Block 8). For the continuous network analysis (dyn-ICA), circuit mean strength and temporal variability were correlated with dual-task benefit for each of the ten circuits. For the discrete-state analysis (HMM), fractional occupancy (FOpost) and number of visits were correlated with dual-task benefit for each of the ten states. Multiple comparisons were controlled separately for each method using the Benjamini–Hochberg false discovery rate (FDR) procedure.

#### 3.2.1 Discovery of behaviorally relevant dyn-ICA circuits

For the continuous network analysis, we correlated two dyn-ICA metrics—mean circuit strength and temporal variability—with dual-task benefit. As shown in Figure 4, two features remained significant after FDR correction across 20 tests (10 circuits × 2 metrics). Greater mean strength of Circuit 4 was positively associated with dual-task benefit (*r* = 0.38, *p* = 0.003, *q* = 0.036), as was greater temporal variability of Circuit 2 (*r* = 0.37, *p* = 0.004, *q* = 0.036). Temporal variability of Circuit 4 showed the strongest remaining positive association (*r* = 0.32, *p* = 0.014), but this effect did not survive FDR correction (*q* = 0.091). Across the other circuits, associations between mean circuit strength and dual-task benefit varied in direction, with several weak negative correlations that did not survive correction. Temporal variability, by contrast, was predominantly positively associated with benefit across circuits, although only Circuit 2 met the FDR-corrected significance threshold. Complete discovery results across all ten circuits are provided in Supplementary Table S2.

#### 3.2.2 Age-group differences in dyn-ICA circuit expression

We next examined whether the two dyn-ICA features associated with dual-task benefit differed between younger and older adults. We then extended the age-group comparison to all ten dyn-ICA circuits to determine whether the effects observed for the behaviorally relevant features were circuit-specific or formed part of a broader age-related pattern of network dynamics (Figures 5–6; Table 3).

**Table 3.** Age-group comparisons of the behaviorally relevant dyn-ICA features.

| Circuit | Metric | Younger adults, Mean (SD) | Older adults, Mean (SD) | t | p value | Cohen's d |
| --- | --- | --- | --- | --- | --- | --- |
| **C2** | Temporal variability | 0.012 (0.004) | 0.018 (0.005) | -4.83 | **1.75x10<sup>-5</sup>** | 1.25 |
| **C4** | Mean strength | -0.029 (0.033) | 0.012 (0.029) | -4.54 | **7.81x10<sup>-5</sup>** | 1.33 |

*Note.* Values are presented as mean (SD). Age groups were compared using Welch's $t$ -tests to accommodate unequal variances. Negative $t$ values indicate higher values in older than in younger adults. Bold type indicates a significant age-group difference at $p < 0.05$ .

As shown in Figure 5, both behaviorally relevant dyn-ICA features differed significantly between age groups. Temporal variability in Circuit 2 was greater in older than in younger adults (older: *M* = 0.018, *SD* = 0.005; younger: *M* = 0.012, *SD* = 0.004; Welch’s *t* = -4.83, *p* < 0.001, Cohen’s *d* = 1.25; Figure 5A, Table 3). Across all participants, greater Circuit 2 temporal variability was associated with a larger dual-task benefit (*r* = 0.37, *p* = 0.004; Figure 5B).

Similarly, the mean strength of Circuit 4 was greater in older than in younger adults (older: *M* = 0.012, *SD* = 0.029; younger: *M* = −0.029, *SD* = 0.033; Welch’s *t* = -4.54, *p* < 0.001, Cohen’s *d* = 1.33; Figure 5C, Table 4). Greater Circuit 4 mean strength was also associated with a larger dual-task benefit across participants (*r* = 0.38, *p* = 0.003; Figure 5D). Thus, the two dyn-ICA features identified through the brain–behavior analysis were both more strongly expressed in older adults and were positively related to behavioral improvement.

**Table 4.** Age-group comparisons of the behaviorally relevant HMM state metrics.

| State | Metric | Younger adults, Mean (SD) | Older adults, Mean (SD) | t | p value | Cohen's d |
| --- | --- | --- | --- | --- | --- | --- |
| **S6** | Number of visits | 0.47 (0.35) | 0.94 (0.36) | -4.80 | **<0.001** | 1.32 |
| **S9** | Fractional occupancy | 0.06 (0.07) | 0.09 (0.08) | -1.73 | 0.090 | 0.44 |

*Note.* Values are presented as mean (SD). Age groups were compared using Welch's *t*-tests to accommodate unequal variances. Negative *t* values indicate higher values in older than in younger adults. Bold type indicates a significant age-group difference at $p < 0.05$ .

To determine whether the age-group differences in Circuits 2 and 4 were selective to the behaviorally relevant features or reflected a broader pattern, we next examined mean strength and temporal variability across all ten dyn-ICA circuits (Figure 6). Older adults exhibited higher mean strength in some circuits (e.g., Circuits 2, 3, 4, 5, and 8) but lower mean strength in others (e.g., Circuits 1, 6, 7, 9, and 10), indicating a circuit-specific reorganization of network expression rather than a global shift (Figure 6A). After applying FDR correction, only Circuits 2, 4, and 8, showed significantly higher mean strength in older adults compared to younger adults. Moreover, these circuits all had positive directional associations with dual-task benefit. In contrast, Circuits 6, 7, and 10, showed significantly higher mean strength in younger adults after FDR correction, and these circuits all had negative directional associations with benefit. Although not every individual circuit–behavior correlation was statistically significant, the direction of association was fully concordant with the direction of age enrichment across all six circuits. Thus, older-enriched mean-strength features were aligned with greater relative performance improvement, whereas younger-enriched features were aligned with smaller relative performance change.

In contrast, temporal variability showed a more consistent age-related pattern. Older adults exhibited greater temporal variability across most dyn-ICA circuits, with the significant differences observed in Circuits 2, 4, 8 and 9 (Figure 6B). Of these, Circuit 2 was the only circuit whose temporal variability was significantly associated with dual-task benefit after correction for multiple comparisons in the discovery analysis. Circuit 4 showed the strongest remaining positive association but this effect did not survive FDR correction. These findings suggest that aging affects dyn-ICA network expression in two distinct ways: a circuit-specific reorganization of mean connectivity strength and a more widespread increase in temporal variability. Successful dual-task adaptation was related to distinct properties of different functional circuits rather than to a uniform increase or decrease in network expression. We further examined the distinct network patterns in these two circuits (Supplementary Figure S5). It revealed that Circuit 2 was centered on visual-attentional sensorimotor reconfiguration, including visual regions and connections with dorsal-attention, sensorimotor, language, cerebellar, and default-mode regions. In contrast, Circuit 4 exhibited a broader integrative network structure including salience, dorsal-attention, frontoparietal, sensorimotor, language, visual, cerebellar, and default-mode regions. Different from Circuit 2, dual-task benefit was associated with its mean strength, not temporal variability. Hence, stronger expression of this circuit, largely in older adults, may facilitate the integration of stimulus detection, attentional control, response selection, and motor execution required for increasingly stable performance.

### 3.3 Discrete network dynamics (HMM)

#### 3.3.1 Discovery of behaviorally relevant HMM states

For the discrete HMM analysis, fractional occupancy (FOpost) and number of visits were correlated with dual-task benefit for each of the ten states. Multiple comparisons were controlled using the Benjamini–Hochberg false discovery rate (FDR) procedure across the 20 HMM state–metric combinations. As shown in Figure 7, two HMM features remained significantly associated with dual-task benefit after FDR correction. Number of visits to State 6 showed the strongest positive correlation (*r* = 0.43, *p* < 0.001, *q* = 0.015), indicating that participants who more frequently visited this state exhibited greater behavioral improvement. In addition, fractional occupancy of State 9 was positively associated with dual-task benefit (*r* = 0.37, *p* = 0.004, *q* = 0.044), suggesting that spending more time in this state also supported dual-task performance. These two behaviorally relevant states were subsequently characterized with respect to age-group differences and their broader network context. Complete discovery results are provided in Supplementary Table S3.

#### 3.3.2 Age-group differences in HMM state expression

We next examined whether the two behaviorally relevant HMM features differed between younger and older adults (Figure 8; Table 4). Number of visits to State 6 was significantly greater in older than in younger adults (older: *M* = 0.94, *SD* = 0.36; younger: *M* = 0.47, *SD* = 0.35; Welch’s *t* = –4.80, *p* < 0.001, Cohen’s *d* = 1.32; Figure 8A). Across all participants, more frequent visits to State 6 were associated with greater dual-task benefit (*r* = 0.43, *p* < 0.001; Figure 8B). In contrast, fractional occupancy of State 9 was only slightly higher in older than in younger adults and did not differ significantly between groups (older: *M* = 0.09, *SD* = 0.08; younger: *M* = 0.06, *SD* = 0.07; Welch’s *t* = –1.73, *p* = 0.090, Cohen’s *d* = 0.44; Figure 8C). Nevertheless, greater fractional occupancy of State 9 was positively associated with dual-task benefit across participants (*r* = 0.37, *p* = 0.004; Figure 8D). Thus, both HMM features predicted behavioral improvement, but only State 6 exhibited a robust age-related increase in expression. The brain connectivity configurations of these two states are shown in Supplementary Figure S6. Both states shared a broadly similar distributed architecture: strong within-network coherence, and coordinated coupling among sensorimotor, salience, and dorsal-attention regions. Compared with State 6, State 9 showed stronger positive coupling across sensorimotor–salience–dorsal-attention regions and more pronounced negative coupling between the default-mode network and salience/dorsal-attention systems.

#### 3.3.3 Network-wide age-group differences in HMM state expression

To determine whether the age-related increase in State 6 reflected a broader redistribution of HMM state expression, we compared fractional occupancy and number of visits across all ten states (Figure 9). Unlike the continuous dyn-ICA metrics, HMM state expression exhibited a state-specific redistribution rather than a global increase. After FDR correction, age-group differences in FOpost were identified for States 6, 7, and 8. Older adults showed higher FOpost for State 6, whereas younger adults showed higher FOpost for States 7 and 8 (Figure 9A). For number of visits, significant age-group differences were observed for States 1, 3, 4, 6, 7, and 8. Older adults visited States 1, 4, and 6 more frequently, whereas younger adults visited States 3, 7, and 8 more frequently (Figure 9B).

To characterize the network configurations underlying these age-related visitation differences, we visualized the state-specific connectivity matrices of the six states whose number of visits differed significantly between age groups after FDR correction (Figure 10). Because a single HMM was estimated across all participants, these matrices represent common state definitions; their organization into older- and younger-preferential columns reflects differences in state visitation rather than connectivity patterns estimated separately for each age group. The figure therefore presents two contrasting age-related visitation profiles characterized by their network configurations and the direction of their associations with dual-task benefit.

The older-enriched profile comprised States 1, 4, and 6, which were visited significantly more frequently by older adults and showed positive associations with relative behavioral improvement (Table 5). This profile was observed alongside older adults’ progressive adaptation from high initial motor RT variability toward their single-motor reference. In contrast, the younger-enriched profile comprised States 3, 7, and 8, which were visited significantly more frequently by younger adults and showed negative associations with relative benefit (Table 5). Because younger adults began near their single-motor reference and rapidly achieved low absolute variability, these negative associations indicate less opportunity for relative improvement rather than poor absolute performance. These findings suggest that younger and older adults preferentially engaged distinct network configurations and that these age-related visitation profiles were aligned with the direction of relative behavioral improvement.

**Table 5.** State-wide age-related expression of HMM states.

| State | Metric | Old – Young difference | Cohen's d | P value | FDR (q) | Direction |
| --- | --- | --- | --- | --- | --- | --- |
| S1 | Visits | 0.241 | 0.59 | 0.026 | 0.043 | Older higher |
| S3 | Visits | -0.344 | -0.81 | 0.008 | 0.024 | Younger higher |
| S4 | Visits | 0.200 | 0.62 | 0.023 | 0.043 | Older higher |
| S6 | FOPost | 0.162 | 1.01 | 0.0002 | 0.0016 | Older higher |
| S6 | Visits | 0.438 | 1.21 | 0.0001 | 0.0010 | Older higher |
| S7 | FOPost | -0.150 | -1.03 | 0.011 | 0.036 | Younger higher |
| S7 | Visits | -0.347 | -0.89 | 0.009 | 0.023 | Younger higher |
| S8 | FOPost | -0.108 | -1.11 | 0.008 | 0.036 | Younger higher |
| S8 | Visits | -0.434 | -1.20 | 0.0038 | 0.019 | Younger higher |

To assess convergence between the two dynamic-connectivity representations, we compared the age-group enrichment and brain–behavior association of dyn-ICA mean-strength features and HMM state-visitation measures (Figure 11). Across both methods, the same directional organization was observed: features preferentially expressed by older adults tended to show positive associations with the dual-task adaptation score, whereas features preferentially expressed by younger adults tended to show negative associations. The strongest FDR-corrected associations within the respective profiles were observed for dyn-ICA Circuit 4 and HMM State 6. Thus, dyn-ICA and HMM showed a consistent directional organization linking age-related feature expression to behavioral adaptation.

**Figure 11.**
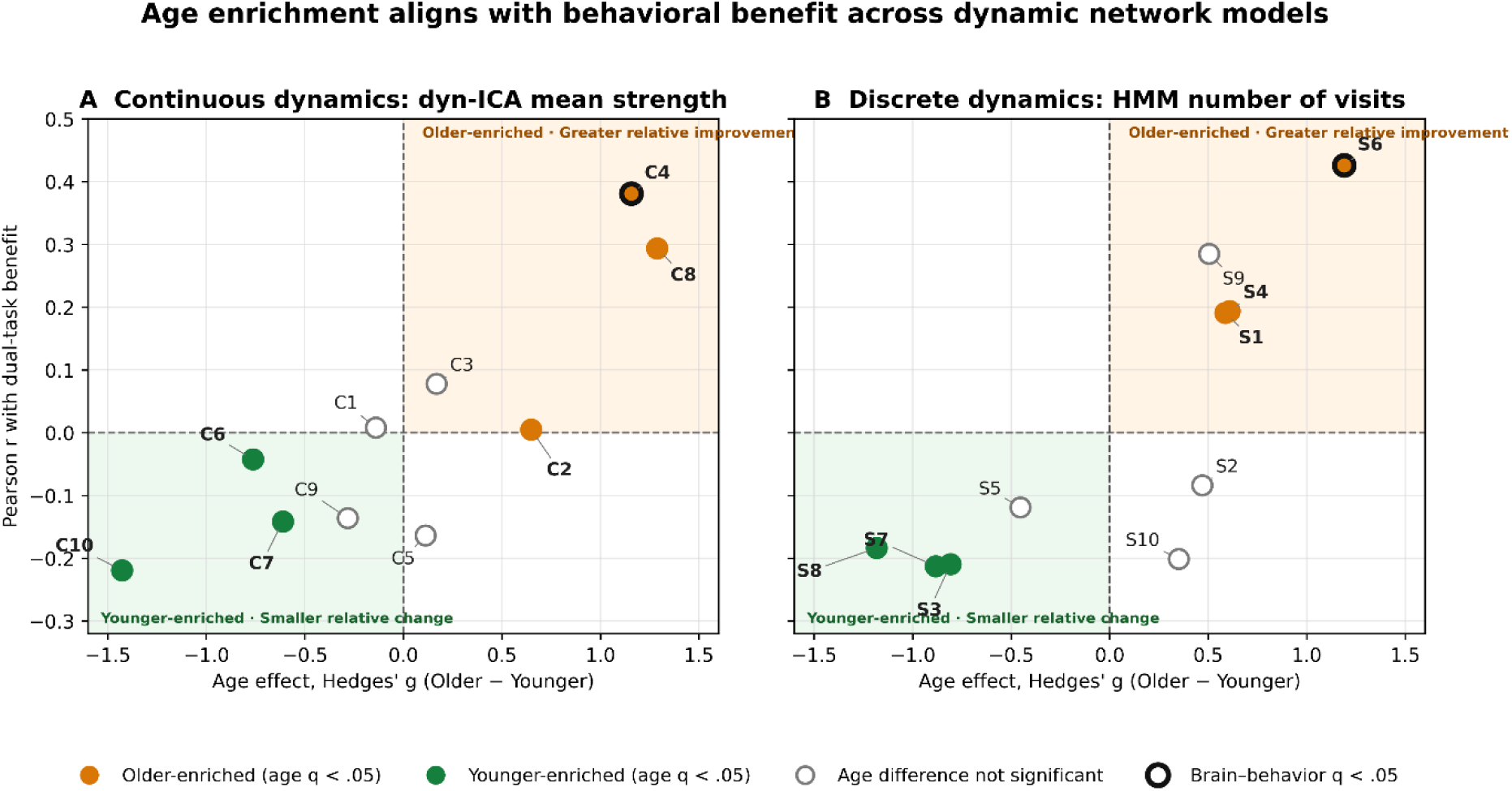
Convergent age–behavior organization across continuous dyn-ICA and discrete HMM representations. The horizontal axis shows the standardized age-group difference (Hedges’ (g), older minus younger), with positive values indicating greater expression in older adults and negative values indicating greater expression in younger adults. The vertical axis shows the Pearson correlation (r) between each dynamic network feature and the dual-task benefit score. (A) Dyn-ICA circuit mean strength. Circuits significantly enriched in older adults (orange) were located in the upper-right quadrant, indicating positive associations with adaptation, whereas circuits significantly enriched in younger adults (green) were located in the lower-left quadrant, indicating negative associations. (B) HMM number of visits. The same directional organization was observed: older-enriched states showed positive associations with adaptation, whereas younger-enriched states showed negative associations. Filled symbols indicate FDR-significant age-group differences; hollow grey symbols indicate features without significant age-group differences. Thick black outlines identify brain–behavior associations that remained significant after FDR correction (dyn-ICA Circuit 4 and HMM State 6). Dashed horizontal and vertical lines mark zero brain–behavior association and zero age-group difference, respectively. Together, the panels demonstrate a convergent directional organization across continuous circuit expression and discrete state visitation.

Together, the dyn-ICA and HMM analyses revealed complementary aspects of age-related brain network dynamics. The continuous dyn-ICA approach identified widespread age-related increases in temporal variability together with circuit-specific changes in mean connectivity strength. In contrast, the discrete HMM analysis revealed a selective redistribution of occupancy and visitation across recurring brain states. Importantly, only a subset of these age-related network changes—Circuit 2 temporal variability, Circuit 4 mean strength, State 6 number of visits, and State 9 fractional occupancy—was significantly associated with dual-task benefit. The convergence of these two analytically distinct approaches revealed a consistent age–behavior organization across continuous circuit expression and discrete state visitation.

## 4. Discussion

We investigated the temporal trajectory of cognitive-motor dual-task performance within a single session in older and younger adults by characterizing their block-wise changes in motor stepping RT variability. By applying continuous (dyn-ICA) and discrete (HMM) models of dynamic brain organization within a common discovery framework, we further identified network features associated with individual differences in behavioral adaptation. Our findings reveal that dual-task performance is not a stationary phenomenon but an adaptive process unfolding over time, and that age-related differences in behavioral trajectories are accompanied by distinct patterns of dynamic network reorganization.

### 4.1 Behavioral adaptation: from static cost to dynamic trajectory

Traditionally, dual-task performance has been interpreted primarily in terms of performance costs relative to single-task behavior (Al-Yahya et al., 2011; Kelly et al., 2010; Plummer & Eskes, 2015). Although this comparison captures the overall interference associated with performing two tasks simultaneously, it does not reveal how performance changes during continued dual-task engagement. Our findings demonstrate that dual-task performance is not temporally stationary but adapts substantially across repeated task blocks. From this dynamic perspective, the initial dual-task cost may represent not the endpoint of cognitive–motor interference, but the starting point of an adaptive process.

Across participants, we observed that motor RT variability was highest during the first dual-task block and progressively decreased over subsequent blocks, reaching its lowest level at Block 6 and remaining relatively stable thereafter. Both age groups showed a substantial early reduction in variability, indicating rapid adaptation during the first few blocks. The progressive reduction in motor RT variability extends the dual-task practice advantage described across longer training periods (Strobach, 2020) to a much shorter timescale, demonstrating that meaningful behavioral adaptation can emerge within a single dual-task session. Although the present design cannot distinguish improved allocation and scheduling from task-specific integration, the association with distributed dynamic network configurations is consistent with increasingly coordinated management of concurrent cognitive and motor demands.

To quantify this within-task performance improvement, we defined *dual-task benefit* as the proportional reduction in motor RT variability in each block relative to the first dual-task block. This measure is conceptually distinct from conventional dual-task cost: whereas cost describes the performance decrement under dual-task relative to single-task conditions (Kelly et al., 2010; Plummer & Eskes, 2015), benefit quantifies improvement over time within the dual-task condition. It therefore characterizes dual-task adaptation as an evolving trajectory rather than as a single comparison between two task conditions (Garner et al., 2020; McPhee et al., 2022).

Despite similar early improvements, the age-group trajectories subsequently diverged. Older adults maintained a relatively large benefit across the later blocks, reaching an approximately 32% reduction in motor RT variability by the end of block (Block 8). In contrast, the benefit in younger adults became smaller during the later phase and was approximately 11% at Block 8. To further understand this group difference, we examined absolute motor RT variability block by block and compared it with each group’s single-motor reference. Younger adults entered the dual-task condition with motor RT variability already close to their single-motor level and showed a rapid reduction below this reference during the early blocks (Blocks 1–2) and remained sub-reference motor variability across most of the dual-task condition. Older adults, by contrast, exhibited markedly elevated variability in the first dual-task exposure, indicating greater initial cognitive–motor interference. However, their variability progressively declined and approached their single-motor reference by approximately Block 6. Thus, the larger relative benefit observed in older adults partly reflects their greater initial impairment—and consequently greater scope for improvement—whereas the smaller late-stage benefit in younger adults should be understood in the context of their comparatively low initial variability.

These findings transform the conventional picture of a static dual-task cost into a temporally evolving process. Initial dual-task interference was followed by progressive stabilization, but the starting points and trajectories differed between the age groups. Younger adults rapidly achieved low motor RT variability, falling below their single-motor reference during the early blocks and remaining below it thereafter. Older adults began with substantially greater instability but progressively recovered single-task-like behavioral stability. The notion of “recovery”, however, requires careful interpretation. Although older adults’ motor RT variability progressively approached their single-motor reference, they did not return to the single-task condition; this stabilization occurred while the concurrent cognitive demands remained present. Their trajectory is therefore more appropriately characterized as adaptation within the dual-task context than as a simple recovery of single-task performance. Moreover, our previous analysis of the same cohort demonstrated significant dual-task costs in motor performance, whereas cognitive performance was relatively preserved (Deng et al., 2026). The observed motor stabilization is therefore unlikely to be attributable solely to abandoning or substantially compromising the concurrent cognitive task. Considered alongside the preferential expression of distributed network configurations, these behavioral findings are consistent with the possibility that older adults used an integrative compensatory process to achieve single-task-like motor stability while continuing to perform the cognitive task. This interpretation remains tentative, however, because cognitive performance was not analyzed block by block; subtle changes in task prioritization over time therefore cannot be ruled out.

This dynamic interpretation complements evidence from dual-task training studies demonstrating functional improvements following repeated practice (Gallou-Guyot et al., 2020; Strobach, 2020; Ye et al., 2024). Although the present within-session changes should not be equated with longer-term training effects, both may reflect adaptive processes unfolding over different timescales. From this perspective, dual-task costs and practice-related benefits are not necessarily opposing phenomena: an initially unstable period of cognitive–motor coordination may be followed by progressive behavioral stabilization with repeated task exposure. Importantly, the pronounced relative benefit in older adults should not be interpreted as evidence of superior overall performance. Rather, it reflects the different starting point and temporal pathway through which stable performance was attained. This behavioral divergence raises a mechanistic question: do younger and older adults achieve stabilization through different patterns of dynamic network organization? We therefore investigated whether the divergent behavioral trajectories were accompanied by age-specific patterns of network dynamics, using dyn-ICA to characterize continuous circuit fluctuations and HMM to identify discrete brain states.

### 4.2 Continuous network dynamics: expression and flexibility in dual-task adaptation

The dyn-ICA analysis identified two complementary continuous network features associated with dual-task benefit after correction for multiple comparisons. Greater mean strength of Circuit 4 was positively associated with behavioral benefit, whereas greater temporal variability of Circuit 2 was also associated with greater benefit. These findings indicate that successful dual-task adaptation was related to distinct properties of different functional circuits.

Circuit 2 was centered on visual regions and included connections with frontoparietal, language, cerebellar, default-mode, dorsal-attention, and sensorimotor regions. Importantly, dual-task benefit was associated with its temporal variability rather than its mean strength, suggesting that adaptation depended more on the dynamic modulation of this circuit than its sustained expression. This interpretation is consistent with evidence that functional connectivity is not stationary but fluctuates among transient configurations over time. Using resting-state fMRI, Allen et al. (2014), demonstrated substantial temporal variation in whole-brain connectivity patterns and proposed that such dynamics reflect flexibility in the coordination of distributed neural systems, with potential relevance to behavioral shifts and adaptive processes. Recent multiscale evidence further indicates that fMRI BOLD variability is spatially heterogeneous and biologically structured, and may support responsivity to changing environmental demands rather than merely representing artefactual noise (Baracchini et al., 2026; Grady & Garrett, 2014). Although these studies examined resting-state connectivity dynamics or regional BOLD-signal variability, whereas our measure captures temporal variability in task-related dyn-ICA circuit expression, they support the broader principle that temporal variability contains biologically meaningful information about dynamic brain organization. In the present dual-task context, greater Circuit 2 variability may reflect a wider dynamic range for adjusting visual processing, frontoparietal motor control, attentional allocation, and sensorimotor coordination as dual-task demands evolve.

The age-group analysis provides additional context for this association. Older adults showed substantially greater temporal variability of Circuit 2 than younger adults, paralleling their larger behavioral improvement from the first to the final dual-task block. However, this difference was embedded within a broader age-related pattern: older adults showed systematically higher temporal variability across all ten dyn-ICA circuits. This finding is broadly consistent with longitudinal evidence that older age is associated with greater variability in functional network organization. Malagurski et al. (2020), for example, found that older age was related to greater global flexibility, as well as greater flexibility within default-mode, frontoparietal-control, and somatomotor networks. Their flexibility measures quantified changes in modular affiliation across four measurement occasions, whereas our measure captured temporal fluctuations in circuit expression within a single dual-task session. Despite these methodological and temporal differences, both findings suggest that aging is accompanied by more variable functional network organization. Importantly, the flexibility identified by Malagurski et al. was not associated with concurrent changes in processing speed or learning performance, indicating that greater network variability is not inherently adaptive. Similarly, Circuit 2 was not unique in showing an age-related increase in our study; rather, it was distinguished by the selective association of its variability with dual-task benefit after correction across all circuits. Thus, the functional significance of elevated variability may depend on the topology of the circuit in which it occurs and its relevance to the sensory, attentional, and motor demands of the task.

Circuit 4 showed a more distributed architecture involving salience, dorsal-attention, frontoparietal, sensorimotor, visual, language, cerebellar, and default-mode regions. Unlike Circuit 2, dual-task benefit was associated with the mean strength of Circuit 4: participants who expressed this configuration more strongly on average showed greater improvement in motor RT stability. This finding is consistent with evidence that task-based whole-brain functional connectivity captures individual differences in RT variability among older adults (Gbadeyan et al., 2022). It also converges with evidence that reorganization of sensorimotor networks accompanies changes in the temporal structure of motor variability during adaptation to altered sensory constraints (Vergotte et al., 2018). Although these studies used different connectivity measures, behavioral outcomes, and task context, they collectively support the broader view that stable motor performance depends on coordinated interactions across distributed functional systems.

This group difference was directionally consistent with both the positive association with dual-task benefit and the greater behavioral benefit observed in older adults. Stronger Circuit 4 expression may therefore represent one network feature involved in coordinating the multiple cognitive and motor processes required for older adults to adapt from their greater initial dual-task instability. It should not, however, be interpreted simply as greater neural efficiency or as direct evidence of compensation (Knights et al., 2025). Stronger expression could reflect adaptive recruitment, greater task demand, or a combination of both. Nevertheless, its association with behavioral improvement suggests that the age-related increase in Circuit 4 strength was functionally relevant rather than merely a nonspecific elevation in network expression.

Taken together, the Circuit 2 and Circuit 4 findings suggest complementary contribution of dynamic flexibility and sustained network expression to dual-task adaptation. The stronger average expression of Circuit 4 may provide a distributed architecture for coordinating attentional control, sensory processing, response selection, and motor execution (Heuninckx et al., 2008). In parallel, greater temporal variability of Circuit 2 may provide the dynamic range needed to adjust visual–attentional and sensorimotor coupling as task demands evolve (Allen et al., 2014; Cohen, 2018). Older adults showed greater mean strength of Circuit 4 and greater temporal variability of Circuit 2, paralleling their pronounced behavioral adaptation from an initially unstable level of performance. These findings suggest that age-related increases in network expression do not necessarily indicate inefficient or maladaptive processing (Goelman et al., 2023; Stojan et al., 2023). When selectively associated with behavioral improvement, they may form part of the dynamic network organization through which older adults adapt to sustained cognitive–motor demands.

Whereas dyn-ICA revealed age-related differences in the magnitude and variability of continuous circuit expression, it does not indicate whether behavioral adaptation also depends on the brain’s occupancy of distinct, recurring network configurations. The HMM analysis addressed this complementary question by characterizing the discrete states visited during dual-task performance and their temporal organization.

### 4.3 Discrete network dynamics: distinct age-related routes to behavioral stability

The HMM analysis identified two behaviorally relevant states after correction for multiple comparisons. More frequent visits to State 6 and greater fractional occupancy of State 9 were associated with greater dual-task benefit. Both states shared a broadly similar distributed architecture: strong within-network coherence, positive coupling within frontoparietal and language systems, and coordinated coupling among sensorimotor, salience, and dorsal-attention regions. Compared with State 6, State 9 showed stronger positive coupling across sensorimotor–salience–dorsal-attention regions and more pronounced negative coupling between the default-mode network and salience/dorsal-attention systems. State 6 appears somewhat less polarized and more diffusely organized. The age-group analysis further revealed a selective redistribution of state expression rather than a uniform increase or decrease across all states.

Older adults visited States 1, 4, and 6 more frequently, collectively forming an older-enriched visitation profile. Note that there was no significant age difference in State 9 FOpost, indicating the similar fractional occupancy expressed by both age groups. This profile was observed alongside their substantial adaptation from initially elevated motor RT variability toward their single-motor reference. State 6 provided the clearest convergence between age and behavior: older adults showed both greater fractional occupancy and more frequent visits to this state, and the number of visits was positively associated with dual-task benefit. State 9, by contrast, was behaviorally relevant but did not differ significantly between age groups, indicating that behavioral relevance and age sensitivity were related but not identical properties. Younger adults visited States 3, 7, and 8 more frequently and also showed greater fractional occupancy of States 7 and 8. The negative directional associations between this younger-enriched profile and relative benefit should not be interpreted as poorer performance. Younger adults began near their single-motor reference and rapidly attained variability at or below this level, leaving less scope for improvement relative to Block 1.

A qualitative comparison of State 6, the state showing the clearest positive association with benefit, and State 7, a younger-enriched state showing a strong negative directional association, illustrated distinct connectivity architectures. State 6 displayed a pronounced block-structured organization, with coherent positive connectivity within several large-scale systems and organized cross-network interactions involving salience, dorsal-attention, frontoparietal, and language regions. By contrast, State 7 exhibited a comparatively finer-grained configuration, with smaller diagonal blocks and less extensive organized structure across networks. These observations suggest that the behavioral relevance of an HMM state may depend not only on how frequently it is visited but also on the network configuration expressed during those visits. However, because network segregation, integration, and modularity were not quantified directly, these differences remain descriptive and should not be interpreted as formal evidence that State 6 was more integrated or State 7 more segregated.

The contrasting visitation profiles are conceptually consistent with previous evidence that younger and older adults differ in how functional network reorganization in response to cognitive and sensorimotor demands. Iordan et al. (2021) found that working-memory training increased network segregation in younger adults, including greater separation of default-mode, frontoparietal/salience, and visual systems. Older adults instead showed diffusely increased between-network connectivity and continued reliance on a more globally integrated organization. Convergent evidence has also been reported within the sensorimotor system. Using resting-state fMRI, Goelman et al. (2023) found relatively direct and predominantly unidirectional sensorimotor pathways in younger adults, whereas older adults exhibited more complex pathways involving mutually coupled higher-order motor regions and inputs originating from visual or insular areas. The authors interpreted this older-adult pattern as reflecting a less segregated and more integrated sensorimotor organization, potentially involving compensatory afferent pathways.

Although these previous studies examined multiday working-memory training or resting-state sensorimotor connectivity, rather than within-session cognitive–motor adaptation, they provide convergent support for an age-dependent distinction in network organization. Younger adults may preferentially engage relatively selective configurations compatible with efficient or increasingly automated performance. Older adults, who experienced greater initial dual-task interference, may instead draw more strongly on distributed configurations involving coordination across multiple functional systems, thereby supporting progressive stabilization under continued dual-task demands. The greater visitation of State 6 among older adults, together with its positive association with behavioral benefit, is compatible with this integrative account. Nevertheless, this interpretation remains provisional because the present analyses did not directly quantify network integration or establish that the observed connectivity architecture caused behavioral stabilization.

Using manifold-based analyses, Mohseni et al. (2026) provide a complementary temporal account of whole-brain reorganization during motor learning. They found that whole-brain neural trajectories were strongly compressed during initial learning, when behavioral errors were greatest, and that these constraints relaxed as performance stabilized. They also linked this evolution to a shift in the dominant contribution of regional activity from sensorimotor toward cognitive-control systems. Although trajectory compression and HMM state expression quantify different aspects of brain dynamics, both approaches indicate that behavioral stabilization is accompanied by systematic reorganization of the available whole-brain dynamic repertoire. In the present study, older adults’ pronounced initial variability may have imposed greater demands on such reorganization, whereas younger adults entered the task closer to a stable regime and therefore required less extensive adjustment.

Taken together, these findings point to two age-related profiles of behavioral stabilization. Older adults followed a distributed adaptation trajectory, preferentially visiting broad network configurations as they progressed from pronounced initial instability toward their single-motor reference. Younger adults followed a more selective maintenance trajectory, beginning closer to this reference and rapidly sustaining low Motor RT variability. Viewed alongside Iordan et al. (2021) and Mohseni et al. (2026), the present findings can be interpreted along two complementary dimensions: age may shape the network configurations used to manage task demands, whereas the stage of adaptation may shape the breadth and flexibility of the available dynamic repertoire. The key observation is therefore not simply that older adults reduced their motor variability, but that they approached single-task-like stability while continuing to perform the concurrent cognitive task. This pattern demonstrates substantial adaptive capacity in older adults and supports a dynamic account of dual-task performance, in which initial interference may be progressively attenuated through changes in the expression and visitation of large-scale network states.

### 4.4 Integrating dyn-ICA and HMM: convergence across complementary representations

Applying dyn-ICA and HMM to the same dataset within a common discovery framework enabled us to compare two complementary representations of brain dynamics. Dyn-ICA captures graded fluctuations in the expression of distributed connectivity circuits, whereas HMM identifies recurring latent configurations and quantifies their temporal visitation. Previous studies have generally applied continuous connectivity methods (e.g., Allen et al., 2014; Chen et al., 2017) or discrete state-based models (Stevner et al., 2019; Taghia et al., 2018; Vidaurre et al., 2017) separately. Their joint application therefore allowed us to examine whether these approaches provided convergent or distinct accounts of dual-task adaptation.

Despite their different mathematical formulations, the two methods revealed a consistent age–behavior organization (Figure 11): features enriched in older adults were generally associated with greater adaptation, whereas features enriched in younger adults showed associations in the opposite direction. This convergence suggests that the age-related organization was not specific to a particular representation of network dynamics. Graded circuit expression and discrete state visitation appear to capture complementary manifestations of a common adaptive architecture. Dyn-ICA showed that adaptation was associated with both the mean strength of Circuit 4 and the temporal variability of Circuit 2, implicating sustained circuit expression alongside flexible modulation. HMM identified complementary associations with State 6 visitation and State 9 fractional occupancy, distinguishing how frequently an adaptive configuration was entered from its overall expression across the task. Together, these findings suggest that behavioral stabilization was related both to continuously varying circuit properties and to the temporal organization of recurring network states—features that could not be fully characterized by either approach alone.

This convergence should not be interpreted as evidence that particular dyn-ICA circuits correspond directly to particular HMM states, or that neural dynamics are fundamentally continuous or discrete. Rather, the two methods provide different lens of a shared dataset and converge at the level of their age–behavior relationships. Their common directional organization strengthens the robustness of the central finding, while their distinct metrics clarify complementary aspects of the adaptation process. Formal quantification of spatial correspondence between circuit and state connectivity configurations will be required to determine whether specific components identified by the two models share network architecture.

### 4.5 Translational implications and limitations

The present findings have implications for how cognitive–motor performance is assessed in healthy aging. Consistent with our previous findings and the broader literature, older adults showed greater overall RT variability than younger adults (Beurskens & Bock, 2012; Bishnoi & Hernandez, 2021; Deng et al., 2026). However, session-average measures obscured substantial within-session adaptation: older adults showed their greatest variability during the initial blocks but progressively approached their single-motor reference as dual-task performance continued. Assessments based on a single block or scan-wide average may therefore conflate transient adjustment demands with persistent performance limitations. Designs incorporating repeated blocks could provide distinguish initial interference, adaptation rate, and stabilized performance more effectively.

Dual-task difficulty may consequently represent not only a means of revealing age-related vulnerability but also a context in which adaptive capacity becomes observable. Despite pronounced initial interference, older adults achieved substantial motor stability while continuing to perform both tasks. This behavioral change was associated with preferential engagement of distributed network configurations, consistent with coordinated adaptation across cognitive and motor systems. Intervention studies provide a rationale for investigating whether such short-term adaptation can be consolidated through repeated practice. For example, Anguera et al. (2022) showed that adaptive cognitive–physical training improved physical fitness and attention-related behavioral and neural outcomes in older adults, with attention benefits maintained one year later. Together with evidence from dual-task training studies (Gallou-Guyot et al., 2020; Schmid, 2025; Strobach, 2020; Y.-R. Yang et al., 2019), these findings suggest that appropriately graded cognitive-motor demands may promote adaptation in healthy aging. The present study does not establish retention or transfer, but it provides a basis for testing whether repeated practice can convert rapidly achieved stabilization into durable functional benefits. The dyn-ICA and HMM findings further illustrate the potential value of temporally sensitive neural markers. Similar late-session behavioral stability was accompanied by distinct age-related profiles of continuous circuit expression and discrete state visitation. Dynamic network features may therefore provide information about how individuals adapt that is not captured by their final performance level alone. Longitudinal studies combining block-resolved behavior with dynamic neuroimaging could test whether these features predict responsiveness to training and help determine the level or type of cognitive–motor challenge most appropriate for a given individual.

These implications should be interpreted within the characteristics of the present sample and design. The study was cross-sectional, involved a relatively small and unequal sample of healthy younger and older adults, and assessed adaptation within a single session. The observed brain–behavior associations are correlational and cannot establish that the identified network configurations caused behavioral stabilization. Participants in older group were relatively healthy older adults with a mean age of 67.6 years. The findings may not generalize to adults over 80 years of age, individuals with multimorbidity or reduced physiological reserve, or populations with cognitive or motor impairment. Finally, cognitive–motor challenge must be matched to individual capacity. Tasks that exceed a person’s functional reserve may increase fall or injury risk, and the present results do not support unsupervised high-challenge activity. Rather, they suggest that appropriately graded and supervised complexity warrants investigation as a component of cognitive–motor interventions. Future longitudinal and interventional studies should establish the populations, task demands, safety conditions, and training doses under which within-session adaptation develops into sustained and transferable benefit.

## 5. Conclusion

Within a single experimental session, healthy younger and older adults achieved behavioral stabilization through distinct adaptation trajectories. Older adults progressed from pronounced initial motor variability toward their single-motor reference while continuing to performance the cognitive-motor dual task. In contrast, younger adults began closer to this reference and maintained comparatively stable performance. Individual differences in adaptation were associated with complementary properties of continuous circuit expression and discrete brain-state visitation. Across both analytical frameworks, older-enriched network configurations were aligned with greater relative behavioral improvement, whereas younger-enriched configurations were associated with smaller relative change from an already more stable starting point. This convergence indicates that age-related adaptation was associated with preferential engagement of specific dynamic network configurations. By integrating dyn-ICA and HMM, the study shows how continuous circuit properties and recurring state dynamics provide complementary understanding of short-term cognitive-motor adaptation. These findings provide a foundation for testing whether repeated practice can consolidate rapidly achieved stabilization into durable and transferable functional benefits.

## 6. Data and code availability

Upon acceptance, all data and analysis code will be uploaded to the Open Science Framework (OSF) and made publicly available.

## 7. Author Contributions

Yan Deng: Conceptualization, Methodology, Software, Formal analysis, Visualization, Investigation, Writing – Original draft preparation Daniel Kristanto: Methodology, Conceptualization, Writing – Reviewing and Editing

## 10. Acknowledgements

We sincerely thank Christiane Thiel and Kayson Fakhar for their substantial contributions to earlier versions of the manuscript, as well as for their valuable input and many insightful discussions that helped shape this work. We further thank Gülsen Yanc and Britta Bruns for their support during MRI scanning. The multiband EPI sequence (CMRR MB-EPI) was kindly provided by the Center for Magnetic Resonance Research (CMRR), University of Minnesota, USA.

