## Supplemental Material for "Continuous and discrete brain dynamics to study behavioral adaptation during cognitive–motor dual-tasking in younger and older adults"

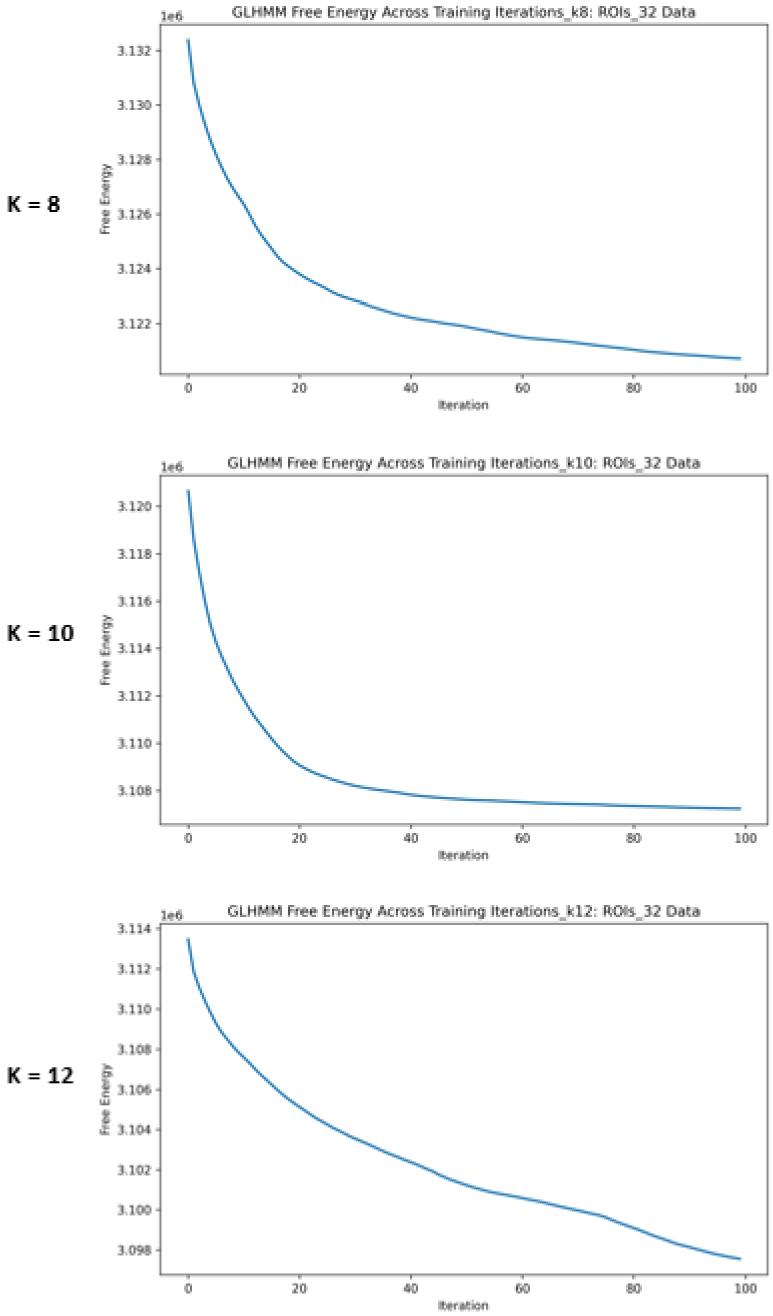


**Supplementary Figure S1. Training convergence of the Gaussian linear hidden Markov models across model orders.** Variational free energy across 100 training iterations is shown for the alternative K=8 solution (top), the primary K=10 solution (middle), and the alternative K=12 solution (bottom), estimated from the 32-region-of-interest dataset. Free energy decreased consistently across iterations for all three model orders. The K=8 and K=10 solutions approached a plateau, whereas the K=12 solution showed a more gradual continued decrease. These trajectories indicate stable model optimization across the model orders examined, although convergence was slower for the higher-order solution.

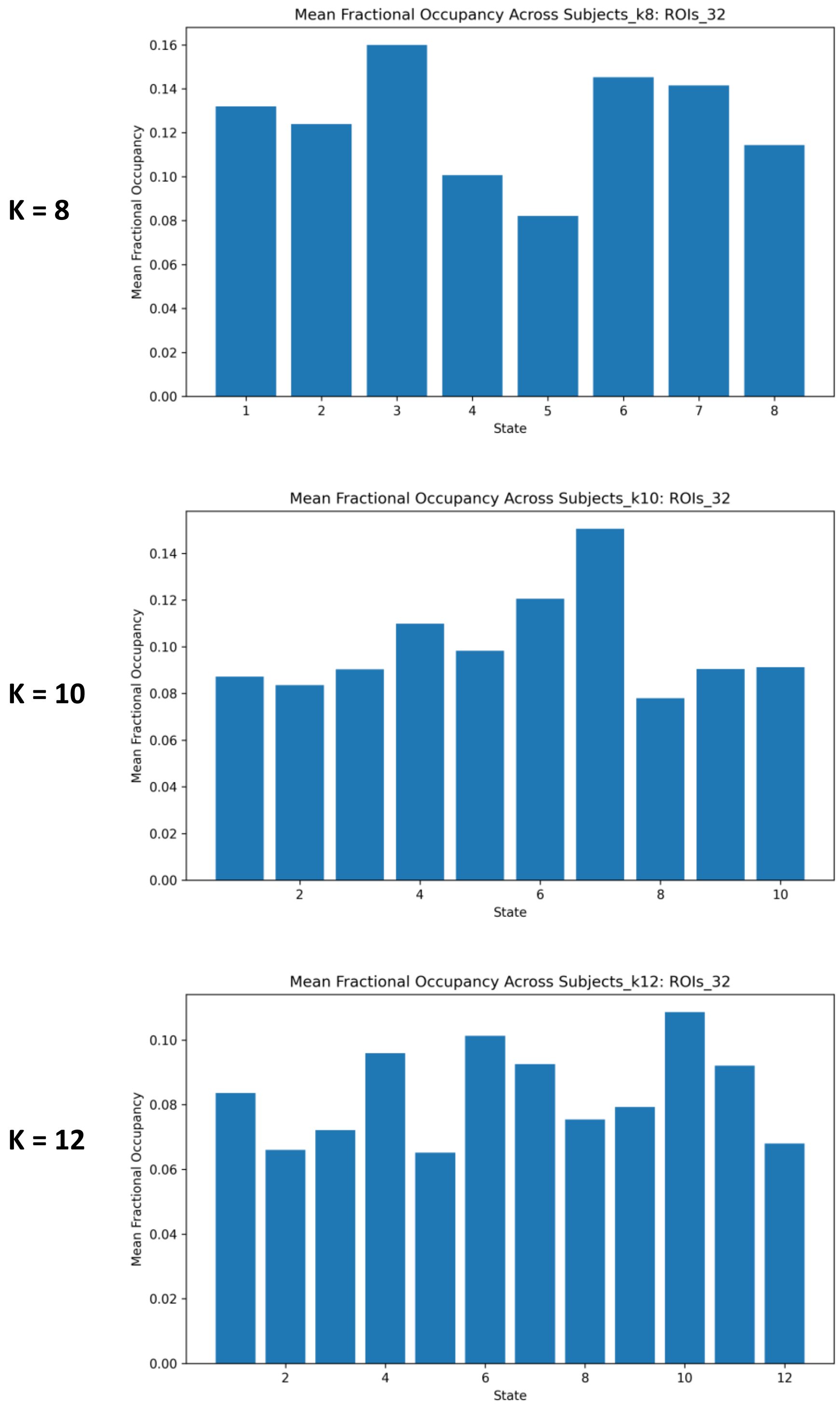


**Supplementary Figure S2. Mean fractional occupancy of HMM states across model orders.** Mean fractional occupancy across participants is shown for the alternative K=8 solution (top), the primary K=10 solution (middle), and the alternative K=12 solution (bottom), estimated from the 32-region-of-interest dataset. All states exhibited non-negligible fractional occupancy at each model order, with no evidence of collapsed or rarely expressed states. Although occupancy was distributed across a larger number of states as K increased, the overall profiles remained relatively balanced, supporting the stability of the HMM solutions across the examined model orders.

K = 8

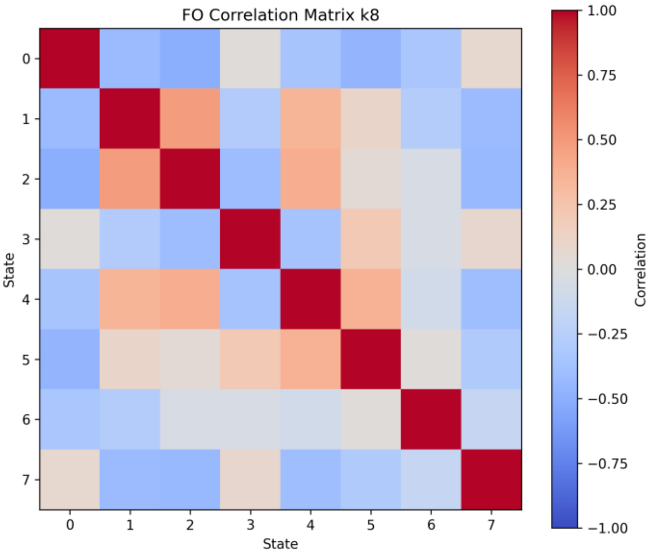
FO Correlation Matrix k8
State
0
1
2
3
4
5
6
7
State
1.00
0.75
0.50
0.25
0.00
−0.25
−0.50
−0.75
−1.00
Correlation

K = 10

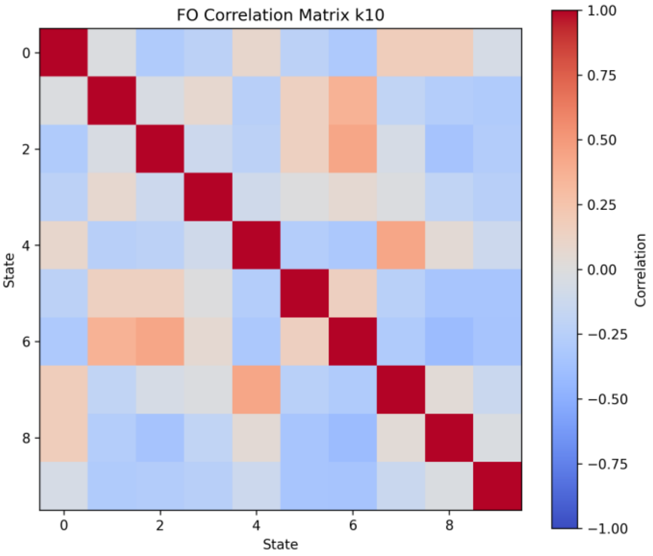
FO Correlation Matrix k10
State
0
2
4
6
8
State
1.00
0.75
0.50
0.25
0.00
−0.25
−0.50
−0.75
−1.00
Correlation

K = 12

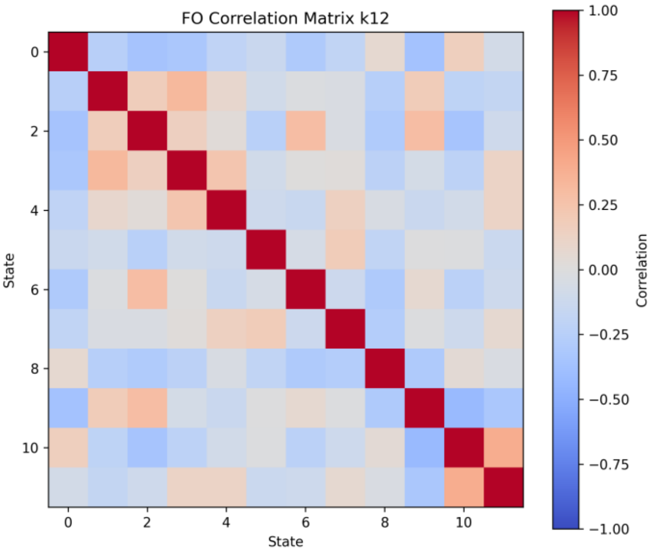
FO Correlation Matrix k12
State
0
2
4
6
8
10
State
1.00
0.75
0.50
0.25
0.00
−0.25
−0.50
−0.75
−1.00
Correlation

**Supplementary Figure S3. Pairwise correlations among HMM-state fractional occupancies across model orders.** Correlation matrices show the across-participant Pearson correlations between the fractional occupancies of all states in the alternative K=8 solution (top), the primary K=10 solution (middle), and the alternative K=12 solution (bottom). Red and blue cells indicate positive and negative correlations, respectively; diagonal values equal 1 by definition. The heterogeneous off-diagonal correlations indicate that the states exhibited distinct occupancy patterns, with no pervasive near-perfect correlations suggesting redundant states. Negative correlations are partly expected because fractional occupancies across states sum to 1 within each participant. States are indexed from 0 to K−1 within each independently estimated solution.

K = 8

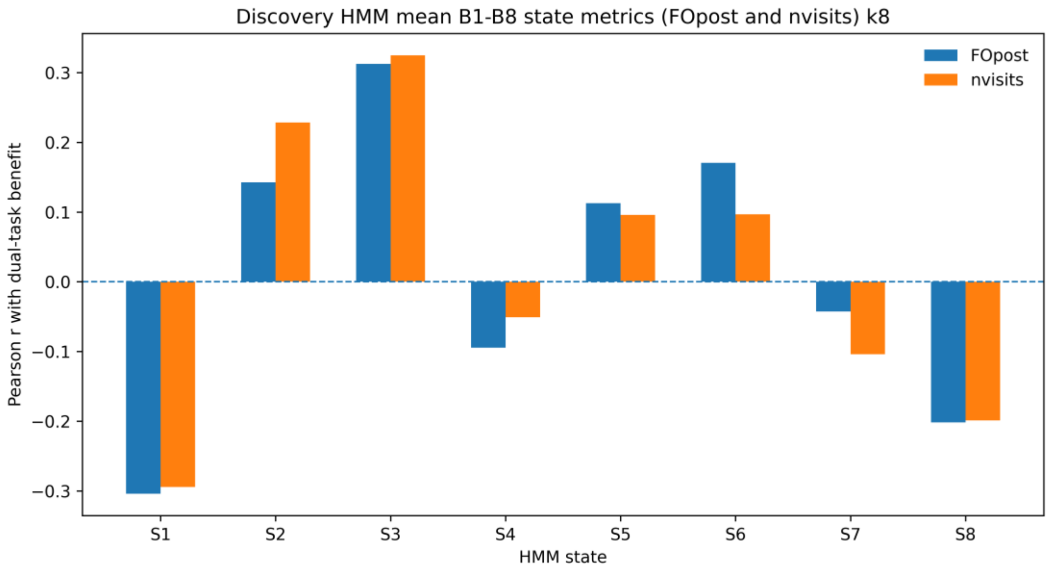
Discovery HMM mean B1-B8 state metrics (FOpost and nvisits) k8
FOpost
nvisits
Pearson r with dual-task benefit
0.3
0.2
0.1
0.0
−0.1
−0.2
−0.3
S1
S2
S3
S4
S5
S6
S7
S8
HMM state

K = 10

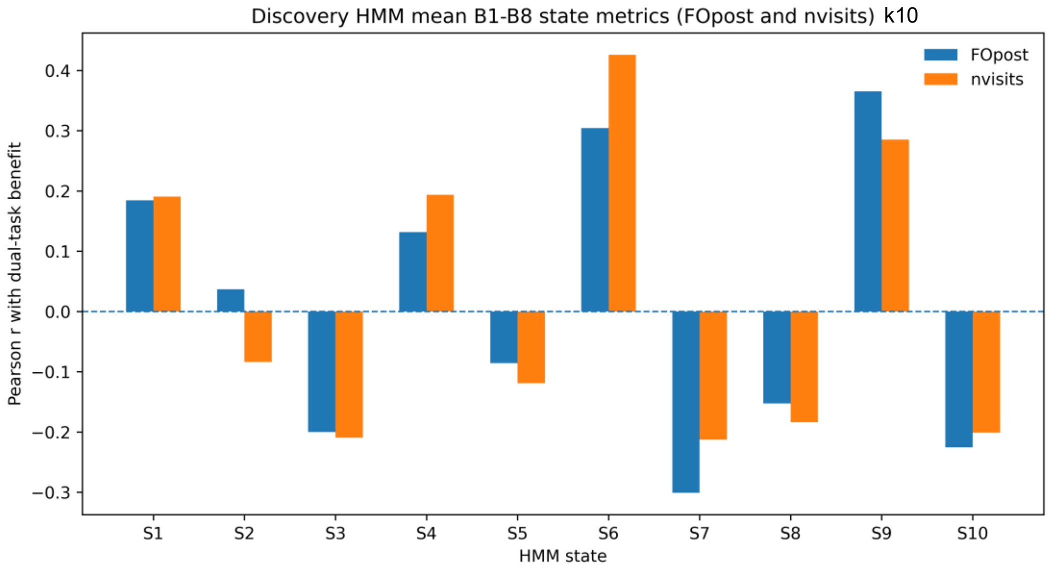
Discovery HMM mean B1-B8 state metrics (FOpost and nvisits) k10
FOpost
nvisits
Pearson r with dual-task benefit
0.4
0.3
0.2
0.1
0.0
−0.1
−0.2
−0.3
S1
S2
S3
S4
S5
S6
S7
S8
S9
S10
HMM state

K = 12

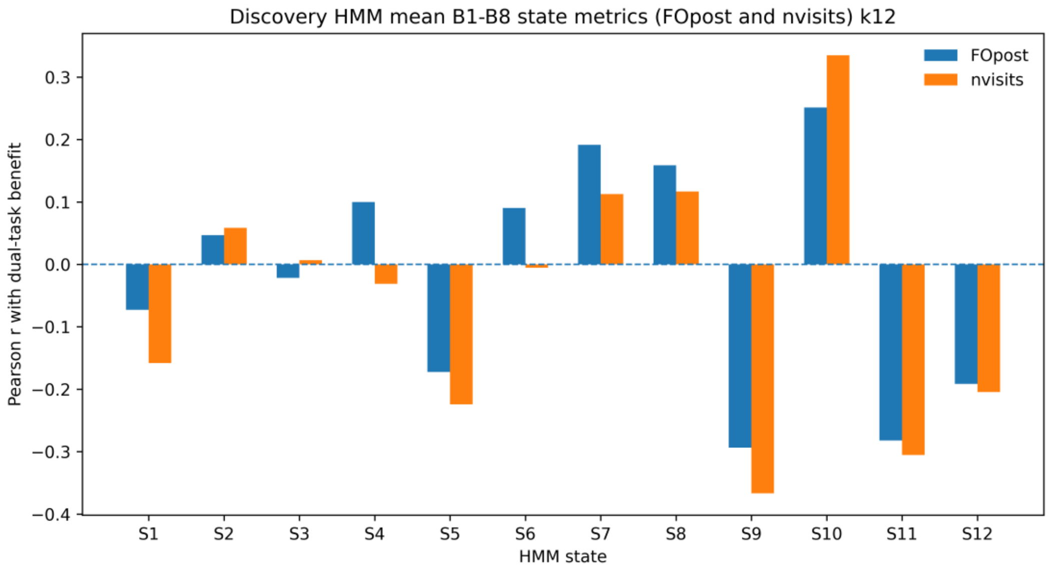
Discovery HMM mean B1-B8 state metrics (FOpost and nvisits) k12
FOpost
nvisits
Pearson r with dual-task benefit
0.3
0.2
0.1
0.0
−0.1
−0.2
−0.3
−0.4
S1
S2
S3
S4
S5
S6
S7
S8
S9
S10
S11
S12
HMM state

**Supplementary Figure S4. Sensitivity of HMM state–behavior associations to model order.** Pearson correlations between dual-task benefit and two HMM state metrics—mean posterior fractional occupancy (FOpost; blue) and mean number of visits (nvisits; orange)—averaged across dual-task Blocks 1–8 are shown for the alternative K=8 solution (top), the primary K=10 (middle), and the alternative K=12 solution (bottom). Across model orders, both positive and negative state–behavior associations were observed, and states showing comparatively strong positive associations with dual-task benefit were consistently identified. This pattern supports the conclusion that behaviorally relevant variation in state expression was not specific to the primary K=10 solution. Bars represent Pearson correlation coefficients. Because each model was estimated independently, state labels are specific to each solution and do not imply one-to-one correspondence across model orders.

**Supplementary Table S1. CONN network ROIs and MNI coordinates**

| Network | ROI | Hemisphere | MNI coordinates (x, y, z) |
|---|---|---|---|
| **Cerebellar** | Anterior | | (0, -63, -30) |
| **Cerebellar** | Posterior | | (0, -79, -32) |
| **Fronto Parietal** | LPFC | L | (-43, 33, 28) |
| **Fronto Parietal** | PPC | L | (-46, -58, 49) |
| **Fronto Parietal** | LPFC | R | (41, 38, 30) |
| **Fronto Parietal** | PPC | R | (52, -52, 45) |
| **Default Mode** | MPFC | | (1, 55, -3) |
| **Default Mode** | LP | L | (-39, -77, 33) |
| **Default Mode** | LP | R | (47, -67, 29) |
| **Default Mode** | PCC | | (1, -61, 38) |
| **SensoriMotor** | Lateral | L | (-55, -12, 29) |
| **SensoriMotor** | Lateral | R | (56, -10, 29) |
| **SensoriMotor** | Superior | | (0, -31, 67) |
| **Dorsal Attention** | FEF | L | (-27, -9, 64) |
| **Dorsal Attention** | FEF | R | (30, -6, 64) |
| **Dorsal Attention** | IPS | L | (-39, -43, 52) |
| **Dorsal Attention** | IPS | R | (39, -42, 54) |
| **Language** | IFG | L | (-51, 26, 2) |
| **Language** | IFG | R | (54, 28, 1) |
| **Language** | pSTG | L | (-57, -47, 15) |
| **Language** | pSTG | R | (59, -42, 13) |
| **Salience** | ACC | | (0, 22, 35) |

| Network | ROI | Hemisphere | MNI coordinates (x, y, z) |
|---|---|---|---|
| **Salience** | AInsula | L | (-44, 13, 1) |
| **Salience** | AInsula | R | (47, 14, 0) |
| **Salience** | RPFC | L | (-32, 45, 27) |
| **Salience** | RPFC | R | (32, 46, 27) |
| **Salience** | SMG | L | (-60, -39, 31) |
| **Salience** | SMG | R | (62, -35, 32) |
| **Visual** | Medial | | (2, -79, 12) |
| **Visual** | Occipital | | (0, -93, -4) |
| **Visual** | Lateral | L | (-37, -79, 10) |
| **Visual** | Lateral | R | (38, -72, 13) |

**Note.** The 32 network ROIs were derived by CONN from ICA of Human Connectome Project data from 497 participants. Network and ROI labels, hemisphere designations, and MNI coordinates were obtained from the accompanying CONN atlas information files.

**Supplementary Table S2. Association between dyn-ICA circuit metrics and dual-task benefit.** Pearson correlation coefficients (r values) are shown for circuit strength and temporal variability across all ten circuits, ranked by absolute correlation strength.

| Circuit | dyn-ICA metric | r | p | FDR-adjusted q | FDR significant |
|---|---|---|---|---|---|
| **Circuit 4** | **Mean strength** | **0.381** | **0.003** | **0.036** | **Yes** |
| **Circuit 2** | **Temporal variability** | **0.373** | **0.004** | **0.036** | **Yes** |
| Circuit 4 | Temporal variability | 0.319 | 0.014 | 0.091 | No |
| Circuit 8 | Mean strength | 0.294 | 0.024 | 0.119 | No |
| Circuit 9 | Temporal variability | 0.227 | 0.084 | 0.287 | No |
| Circuit 10 | Mean strength | -0.219 | 0.095 | 0.287 | No |
| Circuit 3 | Temporal variability | 0.216 | 0.100 | 0.287 | No |
| Circuit 5 | Temporal variability | 0.179 | 0.176 | 0.440 | No |
| Circuit 5 | Mean strength | -0.164 | 0.215 | 0.468 | No |
| Circuit 8 | Temporal variability | 0.157 | 0.236 | 0.468 | No |
| Circuit 6 | Temporal variability | 0.147 | 0.267 | 0.468 | No |
| Circuit 7 | Mean strength | -0.141 | 0.286 | 0.468 | No |
| Circuit 9 | Mean strength | -0.136 | 0.304 | 0.468 | No |
| Circuit 10 | Temporal variability | 0.123 | 0.355 | 0.508 | No |
| Circuit 1 | Temporal variability | 0.080 | 0.545 | 0.697 | No |
| Circuit 3 | Mean strength | 0.078 | 0.558 | 0.697 | No |
| Circuit 7 | Temporal variability | -0.048 | 0.716 | 0.834 | No |
| Circuit 6 | Mean strength | -0.042 | 0.751 | 0.834 | No |
| Circuit 1 | Mean strength | 0.008 | 0.950 | 0.971 | No |
| Circuit 2 | Mean strength | 0.005 | 0.971 | 0.971 | No |

Note. r = Pearson correlation coefficient; FDR = false discovery rate. q values were obtained by applying the Benjamini–Hochberg procedure across all 20 tests (10 circuits × 2 metrics). Exact machine-readable values are retained in the “Exact source data” worksheet.

**Supplementary Table S3. Association between HMM metrics and dual-task benefit.** Pearson correlation coefficients (r values) are shown for fractional occupancy (FOpost), and number of visits (nvisits) across all ten states, ranked by absolute correlation strength.

| State | HMM metric | r | p | FDR-adjusted q | FDR significant |
|---|---|---|---|---|---|
| **State 6** | **Number of visits** | **0.426** | **< .001** | **0.015** | **Yes** |
| **State 9** | **Fractional occupancy** | **0.365** | **0.004** | **0.044** | **Yes** |
| State 6 | Fractional occupancy | 0.304 | 0.019 | 0.103 | No |
| State 7 | Fractional occupancy | -0.301 | 0.021 | 0.103 | No |
| State 9 | Number of visits | 0.285 | 0.029 | 0.114 | No |
| State 10 | Fractional occupancy | -0.225 | 0.086 | 0.234 | No |
| State 7 | Number of visits | -0.213 | 0.106 | 0.234 | No |
| State 3 | Number of visits | -0.209 | 0.111 | 0.234 | No |
| State 10 | Number of visits | -0.201 | 0.126 | 0.234 | No |
| State 3 | Fractional occupancy | -0.200 | 0.129 | 0.234 | No |
| State 4 | Number of visits | 0.194 | 0.141 | 0.234 | No |
| State 1 | Number of visits | 0.191 | 0.148 | 0.234 | No |
| State 1 | Fractional occupancy | 0.184 | 0.162 | 0.234 | No |
| State 8 | Number of visits | -0.184 | 0.164 | 0.234 | No |
| State 8 | Fractional occupancy | -0.153 | 0.248 | 0.331 | No |
| State 4 | Fractional occupancy | 0.132 | 0.320 | 0.400 | No |
| State 5 | Number of visits | -0.119 | 0.369 | 0.434 | No |
| State 5 | Fractional occupancy | -0.086 | 0.518 | 0.555 | No |
| State 2 | Number of visits | -0.084 | 0.528 | 0.555 | No |
| State 2 | Fractional occupancy | 0.037 | 0.782 | 0.782 | No |

Note. r = Pearson correlation coefficient; FDR = false discovery rate. q values were obtained by applying the Benjamini–Hochberg procedure across all 20 tests (10 states × 2 metrics).

### Circuit 2 functional connectivity

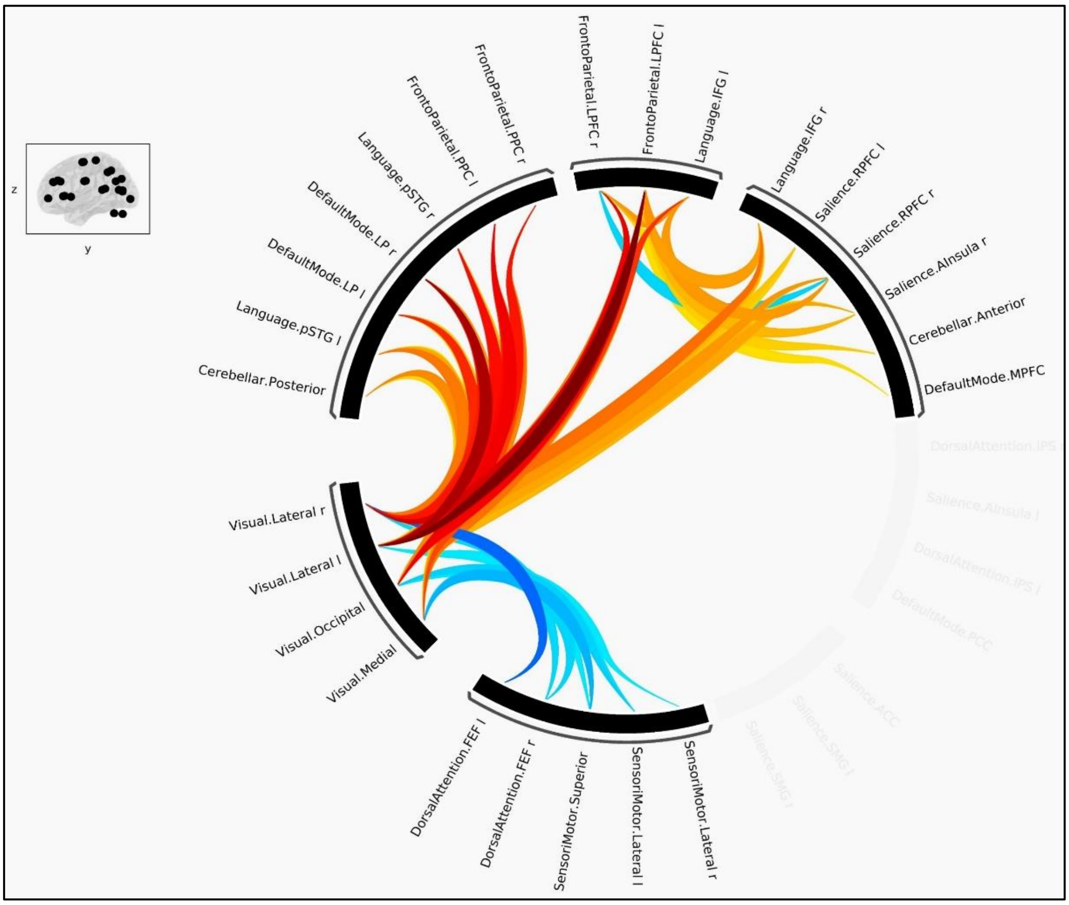


### Circuit 4 functional connectivity

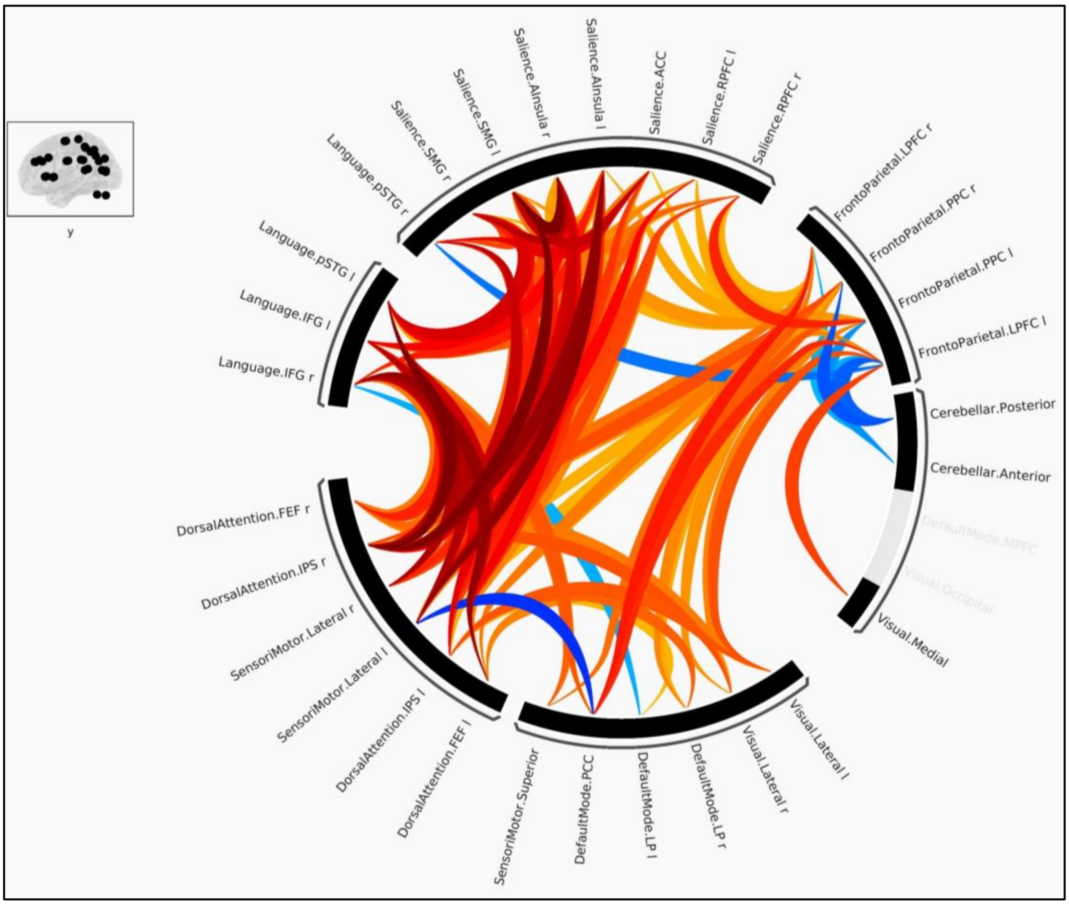

**Supplementary Figure S5. Functional-connectivity configurations of behaviorally relevant dyn-ICA circuits.** Chord diagrams show the connectivity configurations of (A) Circuit 2 and (B) Circuit 4 during the cognitive–motor dual-task condition. Connections are arranged according to the 32 ROIs of the CONN network atlas. Edge colors represent the sign and magnitude of the corresponding connectivity statistic, with warmer colors indicating positive values and cooler colors indicating negative values. Line width reflects the magnitude of the effect.

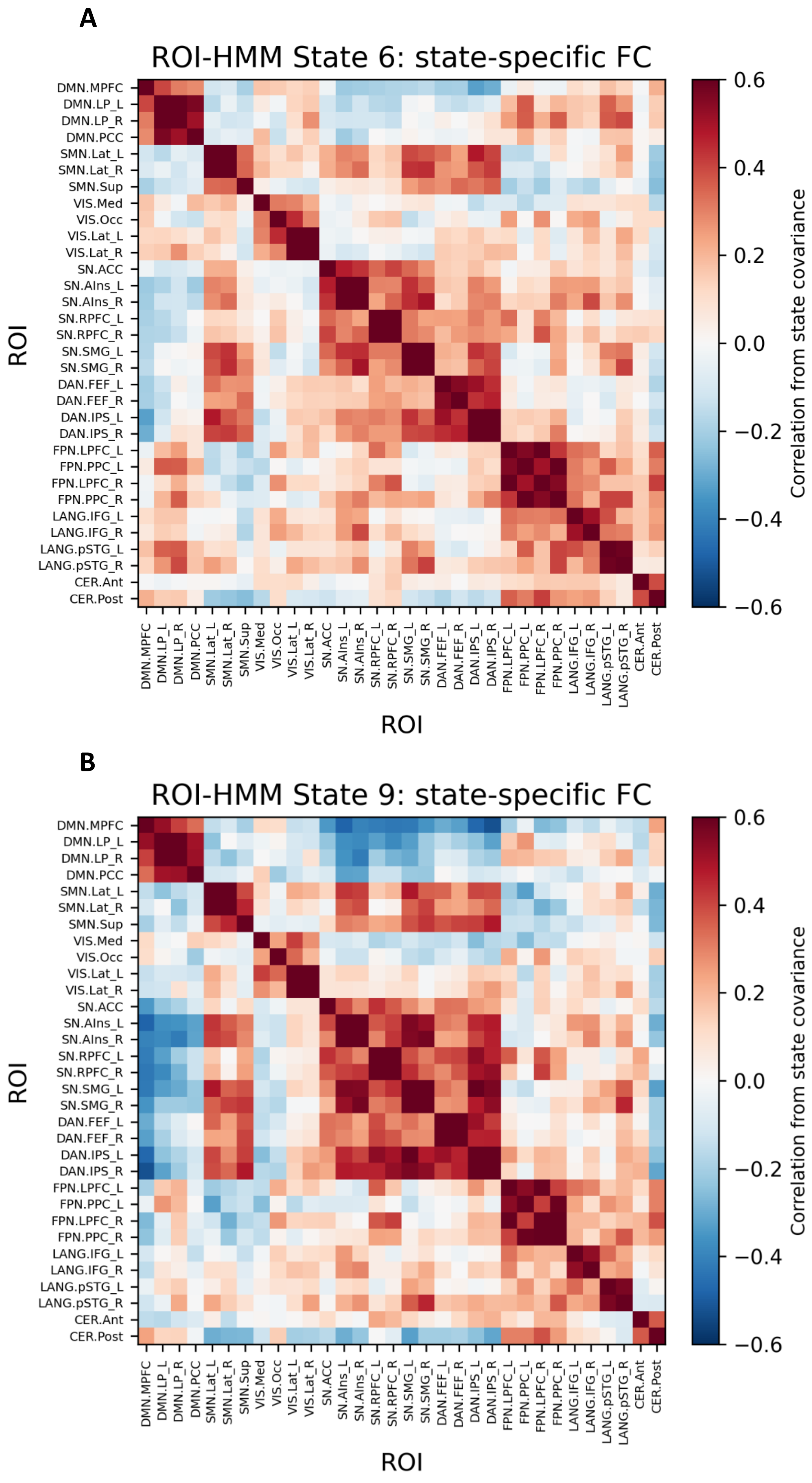
A
ROI-HMM State 6: state-specific FC
ROI
Correlation from state covariance
0.6
0.4
0.2
0.0
−0.2
−0.4
−0.6
B
ROI-HMM State 9: state-specific FC
ROI
Correlation from state covariance
DMN.MPFC
DMN.LP_L
DMN.LP_R
DMN.PCC
SMN.Lat_L
SMN.Lat_R
SMN.Sup
VIS.Med
VIS.Occ
VIS.Lat_L
VIS.Lat_R
SN.ACC
SN.AIns_L
SN.AIns_R
SN.RPFC_L
SN.RPFC_R
SN.SMG_L
SN.SMG_R
DAN.FEF_L
DAN.FEF_R
DAN.IPS_L
DAN.IPS_R
FPN.LPFC_L
FPN.PPC_L
FPN.LPFC_R
FPN.PPC_R
LANG.IFG_L
LANG.IFG_R
LANG.pSTG_L
LANG.pSTG_R
CER.Ant
CER.Post

**Supplementary Figure S6. Connectivity configurations of two behaviorally relevant HMM states.** (a) HMM State 6 and (B) HMM State 9. Both states shared a broadly similar distributed architecture: strong within-network coherence, positive coupling within frontoparietal and language systems, and coordinated coupling among sensorimotor, salience, and dorsal-attention regions. Compared with State 6, State 9 showed stronger positive coupling across sensorimotor–salience–dorsal-attention regions and more pronounced negative coupling between the default-mode network and salience/dorsal-attention systems.
